# A new cancer progression model: from synthetic tumors to real data and back

**DOI:** 10.64898/2026.02.06.704299

**Authors:** Daniela Volpatto, Sandro Gepiro Contaldo, Simone Pernice, Marco Beccuti, Francesca Cordero, Roberta Sirovich

**Affiliations:** Department of Computer Science, University of Torino, Turin, Italy; Department of Mathematics G. Peano, University of Torino, Turin, Italy

## Abstract

Intratumor heterogeneity (ITH) arises from the combined effects of genetic alterations, clonal interactions, and environmental constraints, and plays a central role in therapeutic resistance and disease progression. While ITH has been extensively documented in empirical tumor data, the scientific debate regarding the biological mechanisms underlying this heterogeneity remains complex, highlighting the need for cancer evolution models that are sufficiently flexible and sophisticated to reproduce the observed behaviors and to give insights on the unobserved ones. Here, we present a stochastic modelling framework for tumor evolution that integrates genotypic inheritance with phenotype driven functional traits and resource mediated competition. Mutational events are associated with functional capabilities such as altered proliferation, increased mutation rates, limit evasion potential or enhanced control over shared resources, allowing multiple genotypes to converge on similar phenotypes. The model explicitly tracks subclonal lineages while incorporating environmental constraints that modulate growth and competition.The framework is defined through a mathematically rigorous construction and is accompanied by an efficient simulation algorithm. To facilitate exploration and reproducibility, we provide an open-source graphical user interface that allows users to configure model parameters, run simulations, and inspect clonal genealogies and population dynamics without requiring direct interaction with the underlying code. Using this model, we illustrate how ecological feedbacks can shape clonal dynamics over time, supporting an interpretation in which early tumor growth is dominated by stochastic expansion, while later evolution increasingly reflects selection for traits that alleviate environmental constraints. Rather than constituting a new evolutionary paradigm, this behaviour demonstrates how well-documented biological patterns can emerge naturally from a unified stochastic and ecological description. Overall, our approach offers a flexible and extensible platform for investigating how chance, functional traits, and environmental interactions jointly govern tumor heterogeneity.

**Author summary:** Not all cancerous cells are created equal: inside the same tumor, different populations of cells exist at the same time, fighting for the same resources and influencing the way the disease evolves and reacts to treatments. These groups of cells have different behaviour and abilities thanks to different genetic mutations, which might give them an advantage or bring their population to disappearance. We have built a mathematical model that mimics the evolution of a tumor over time, simulating a competition between its different populations of cells. Our simulated experiments show that tumors evolve in two distinct phases: at first, cells that grow and divide more quickly have an advantage. Once the space and nutrients are limited, cells that can survive with fewer resources have an advantage and can potentially take over the race. We use these simulations to argue that the evolution of a tumor doesn’t depend on the shape of the space it expands in, but rather on the availability of nutrients.

## Introduction

Cancer is a profoundly heterogeneous disease, characterized by the coexistence of multiple subpopulations of cells within a single tumor, each potentially carrying distinct genetic, epigenetic, and phenotypic features. This phenomenon, known as *intratumor heterogeneity*, poses significant challenges for diagnosis, prognosis, and treatment [1–4]. Understanding the dynamics and structure of the cancer heterogeneity is crucial to improving patient outcomes. In particular, intratumor heterogeneity (ITH) drives therapeutic resistance, makes it difficult to detect cancer driving mechanisms and often underlies relapse after initially successful treatment [5–8].

Traditional approaches to cancer research, largely grounded in molecular biology and genomics, have illuminated many aspects of tumor heterogeneity enabling detailed characterization of tumor cell populations, their genetic mutations, metabolic phenotypes, and microenvironmental features. However, these experimental methods often struggle in capturing the dynamics of tumor evolution, especially the temporal dimension that traces how subclonal populations emerge, expand, and compete over time [7,9,10]. Reconstructing the evolutionary history of a tumor remains one of the key challenges in this field indeed. In human patients, serial biopsies across disease progression are rarely feasible due to ethical and practical constraints; as a result, most evolutionary inferences must be drawn from single timepoint data, leveraging the fact that the genetic diversity within a tumor encodes a molecular archive of past mutational events.

This challenge has given rise to the interdisciplinary field of *tumor evolution*, which draws on concepts from evolutionary biology, ecology, and population genetics to study how tumor cells adapt to selective pressures over time [1, 5, 11]. Within this framework, mathematical modeling has emerged as a powerful complementary approach [12, 13]. Mathematical models allow researchers to go beyond observational data and test hypotheses about the underlying processes that shape heterogeneity. They enable simulations of evolutionary scenarios and predictions of clonal behaviour under different selective pressures by formalizing assumptions and linking them to measurable outcomes.

According to [10], theories regarding evolutionary dynamics of tumors proposed in the past decade can be conceptualized through four main models: *Linear Evolution*, according to which new driver mutations sequentially outcompete previous clones, resulting in a dominant clonal population [14, 15]; *Branching Evolution* where multiple subclones arise from a common ancestor and evolve in parallel, creating a highly branched phylogenetic tree (this model is supported by numerous genomic studies and explains the persistent coexistence of multiple subclones within tumors) [16–21]; *Neutral evolution* posits that many mutations accumulate without strong selective pressure, resulting in a mixture of subclones with similar fitness and a characteristic allele frequency spectra [22, 23]; in *Punctuated evolution*, tumors undergo a bursts of rapid genomic change, often associated with catastrophic events such as chromosomal rearrangements, at a very early stage of the disease. After this punctuated event, one or a few dominant clones stably expand to form the tumor mass [24]. Each model has different implications for diagnosis, prognosis, and treatment. Importantly, evidence suggests that they are not mutually exclusive; different regions of the same tumor, or different classes of mutations, may follow distinct evolutionary trajectories and the very same tumor might undergo modes in different stages.

Mathematical and computational models have been central to formalizing these concepts [24–26]. Mathematical models of tumor evolution broadly fall into two classes: well-mixed formulations and spatially structured formulations, each of which can be implemented through deterministic or stochastic approaches [12, 13, 27]. Deterministic models (typically ODE, PDE, or hybrid systems) offer biological detail and can capture growth–consumption dynamics, microenvironmental interactions, and spatial diffusion [28–32]. Their main limitation is that they describe average trajectories, which makes them suboptimal for representing intratumor heterogeneity, a phenomenon fundamentally driven by rare, stochastic events [27]. Agent-based models can also capture spatial heterogeneity at single-cell or component resolution, while accounting for stochasticity. However, they are often computationally intensive and allow limited analysis of mathematical properties, being mainly supported by simulations [27]. This leads back to the classical but powerful tools of stochastic processes for population evolution. Two main instruments have been exploited within this area: branching processes, which generate exponential growth by construction, and Moran (or modified Moran) processes, which impose a fixed population size. Attempts to introduce density dependence exist [33], but they remain restricted to a few population types or biologically narrow scenarios. Furthermore, in previously developed models, selective advantage is typically encoded only as a proliferative boost, despite the fact that fitness in tumors depends heavily on resource availability, tolerance to deprivation, and competitive or cooperative interactions among clones [12, 15, 33–36]. Evolutionary game-theoretic frameworks also have explored interaction-driven selection, though they tend to focus on equilibrium coexistence rather than full evolutionary trajectories [12, 13]. Phenotype-driven models have been proposed, but they typically target specific mechanisms and capture only a narrow range of phenotypic classes, while ignoring genomic structure [12, 37].

We propose here a new model for tumor evolution that integrates resource constraints, cellular interactions, and phenotypic rules, without losing the ability to represent genotypes and subclonal genealogies, by adopting a broad definition of clones and functional mutational events. Recent spatial stochastic models [38, 39] move in a similar direction, but often rely on the assumption that spatial segregation is the primary driver of evolutionary mode. We argue that equivalent patterns can arise from interaction-mediated heterogeneity, even in the absence of sharp spatial segregation. Our goal is to provide a flexible, event-based stochastic framework that unifies evolutionary and ecological processes within a single architecture. It supports diverse functional events beyond driver and passenger schemas, incorporates carrying capacity to reproduce both early and late growth regimes, explicitly tracks subclonal genealogies, and is modular, making it straightforward to integrate emerging biological insights. In doing so, we bring together, to the best of our knowledge, aspects that have previously been studied only in isolation within a single coherent framework.

## Materials and methods

### The cancer progression model

Assuming that a tumor originates from a single cell that, as a result of one or more mutational events (genetic or epigenetic), gains the ability to proliferate uncontrollably, bypassing cellular growth regulation mechanisms and invading surrounding tissue, we model the evolution of its subclones as a stochastic process built over a rooted ordered tree. Over time, mutational events can accumulate, creating a heterogeneous set of cellular populations, each characterized by specific genetic traits that influence their ability to survive, replicate, resist therapies and interact with the surrounding environment.

Informally, the model we are going to describe can be thought of as a cell duplicating and dying at certain birth and death rates (*a* and *b*); sometimes, at a certain rate *µ*, the perfect duplication could fail, giving rise to a new subclone carrying an additional mutational event. In this framework, mutational events can encompass any genetic structural variation or epigenetic alteration. The only requirement is that these events are irreversible: once a cell acquires a mutation, it will retain it permanently and pass it on to all its descendant cells. A population (or clone) consists of cells that share a unique, defining genotype, which is stored with a unique identifier.

We aim to describe the size of each population of cells as it changes over time, hence we will build a process *X*_*u*_(*t*), where *u* will indicate the identifier of the population. Since only the original population starts its course at time 0, for all others, we will have a random time of appearance *σ*_*u*_. Furthermore, we will need a structure to record the genetic relationships between populations, which will be constructed by exploiting *rooted ordered trees*, a mathematical structure that encodes the phylogenetic tree of a whole population.

To each population will be associated a *phenotype*. By phenotype, we refer to the set of functional traits that define a population with a specific genotype. A single phenotype can be determined by different genotypes, as different mutations can lead to similar functional outcomes. The phenotype of a population will determine the rules governing its birth-and-death dynamics, the likelihood of new mutations arising, and the cells’ abilities to acquire additional resources and bypass the physiological constraints of a healthy organism.

The overall tumor model will result from the combination of these elements working at different levels. On one hand, we model the genetic makeup of cell populations that descend from a single ancestor and are organized within a tree of inheritance and parental relationships, which drive the evolutionary trajectory of the tumor. On the other hand, the model incorporates the mapping of each genotype to its observable biological effects-the phenotype-which contributes to tumor progression by influencing the final cellular composition. A graphical representation of the different components of the model is shown in Fig. 1.

**Fig 1.**
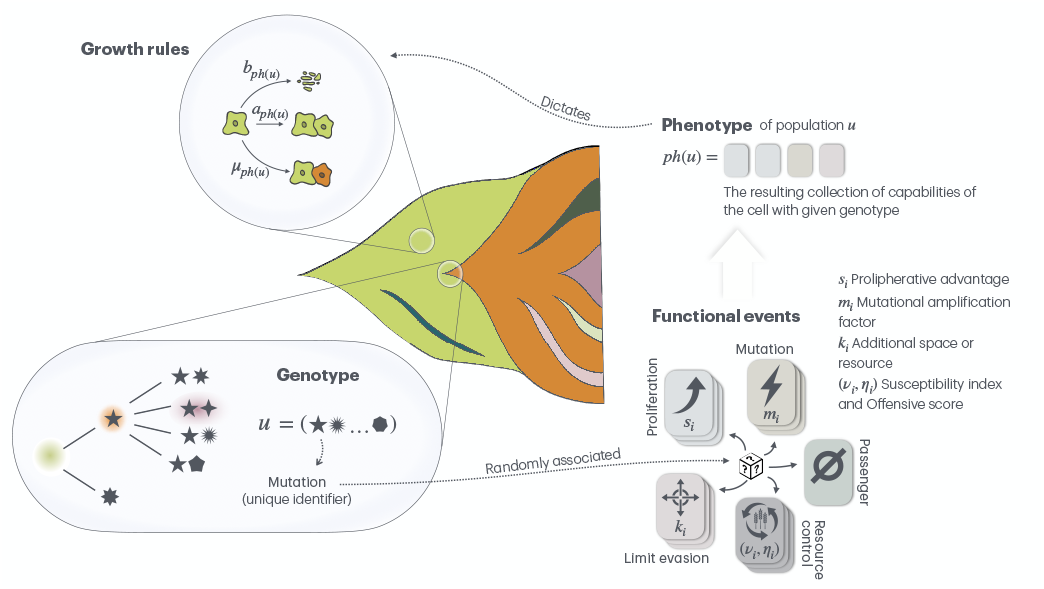
Visual representation of the cancer progression model.

In the next Sections we will formally introduce each of these elements.

#### Genotype and Rooted Ordered trees

The data structure we will use for the description of the phylogenetic evolution of a tumor is the rooted tree. Rooted trees are collections of elements of the set 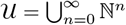 made of the finite sequences of natural numbers ℕ, with ℕ^0^ = ∅. On the generic element *u* = (*u*_1_,..., *u*_*n*_) ∈ 𝒰 we can define |*u*|:= *n* the generation of *u, p*(*u*)= (*u*_1_,..., *u*_*n*−1_) the parent of *u* and we will denote as *u*|_*j*_:= (*u*_1_,…, *u*_*j*_) the restriction up to generation *j* of the element *u*. A *locally finite ordered rooted tree* (LFORT) *τ* is a subset of 𝒰 satisfying:

1. Ø ∈ *τ*
2. *u* ∈ *τ* ⇒ the parent *p*(*u*)= (*u*_1_,..., *u*_*n*−1_) ∈ *τ*
3. ∀*u* ∈ *τ* ∃*A*_*u*_ ∈ ℕ∪0: *uj* = (*u*_1_,..., *u*_*n*_, *j*) ∈ *τ* ⇔ *j* ≤ *A*_*u*_.

The number *A*_*u*_ is the number of children of *u*. We will denote with 𝒯 the collection of all LFORTs. An example of LFORT is illustrated in Fig. S1.1.

From a tumor modeling perspective, a tree is the genetic lineage from the common ancestor placed as the root. Each element *u* of a tree can be interpreted as a genotype, i. e. a *mutational profile*. A new genotype *v* branches from *u* at the (*n* + 1)-th generation when an additional mutational event arises, with parent equal to *u*, i. e. *p*(*v*)= *v*|_*n*_= *u*.

The tree can be enriched with extra structure: for each *u* ∈ *τ* we associate *σ*_*u*_ ≥ 0, the birth time of *u*, and (*X*_*u*_(*t*),*t* ≥ *σ*_*u*_) the number of cells, with genotype *u*, at time *t*. Formally, borrowing the nomenclature from the phylogenetic literature, we can define the following objects: the population *X*_*u*_ with *u* = (*u*_1_,..., *u*_*n*_) ∈ *τ*, the subpopulation (or daughter population) of *X*_*u*_, i.e. *X*_*uj*_ for some *j* ∈ ℕ with *uj* = (*u*_1_,..., *u*_*n*_, *j*), the descendant population *X*_*v*_, with *v* = (*v*_1_,..., *v*_*m*_) ∈ *τ, m* ≥ *n* and such that *v*|_*n*_= *u*, the ancestor population *X*_*v*_ with *v* = (*v*_1_,..., *v*_*j*_) ∈ *τ, j* ≤ *n* and such that *u*|_*j*_= *v*. We will identify a clone as the set of all the descendant populations of *u* and a subclone as a clone with root *v* that has *u* as ancestor.

#### The phenotype as the genotype-associated functional capabilities of a cell

The mutational events may or may not induce changes in the functional traits of the cell. We will refer to the collection of eventual alterations of the biological processes of a cell with a given genotype *u* as the *phenotype*, meaning with that the observable effect that the mutational events have on it. We are following the framework developed by D. Hanahan and R. A. Weinberg, first outlined in “The Hallmarks of Cancer” [40] and later expanded in [41] and [42], that organizes the diverse capabilities and enabling characteristics acquired during the multistep development of human tumors. They claim:

> “We foresee cancer research developing into a logical science, where the complexities of the disease, described in the laboratory and clinic, will become understandable in terms of a small number of underlying principles. Some of these principles are even now in the midst of being codified. […] We suggest that most if not all cancers have acquired the same set of functional capabilities during their development, albeit through various mechanistic strategies.”

These hallmark capabilities - each supported by specific mutations - describe how cancer cells diverge from normal cellular functions to support unchecked growth and adaptability within the body. We developed the phenotype part of our model using the conceptual framework provided by Hanahan et al. [40–42] with the aim of including recognised functional characteristics of human tumours and mapping them into a clear mathematical description. We propose that, from a mathematical modeling perspective, the functional capabilities can be further merged into five primary mechanisms that affect the evolutionary processes of the cells:

1. **Deregulation of the proliferation program** (dividing faster/dying slower). We map into this class all the functional capabilities that have an effect on the replication process of the cells: we may therefore include here the acquired abilities in sustaining the proliferative signaling, in evading growth suppressors, in deregulating the cellular metabolism, in resisting cells death, in enabling the replicative immortality and in avoiding the immune destruction. The simplified functional effect is a a boost in the growth of the cell by either diminishing the expected time required before a cell encounters duplication -a progression through the cell cycle- and by increasing the replicative potential -immortality-, or enlarging lifespan -circumvention of the apoptotic program-. The homeostasis of cell number and the maintenance of normal tissue architecture and function is lost and a surplus in the number of births compared to the number of deaths is observed.
2. **Mutation burden augmentation** (mutating more often). Whenever the DNA is duplicated, there is a possibility of running into an error: this can be measured in terms of number of errors per cell division divided the number of base-pairs in order to obtain a standard mutation rate. There is evidence that the acquisition of the hallmarks of cancer is made possible by several enabling characteristics, among which the most prominent is the development of genomic instability that increases the mutation rate on tumor cells, as the succession of the alterations in the genomes of neoplastic cells results in the acquisition of function-altering mutations which enable the development of different capabilities.
3. **Limit evasion** (potential for expansion over defined physiological limits). Tumors are located within a body, hence they are subject to the physical constraints and to the limitation of the available resources. The infrastructure of the tissues in which cancer develops are built to bear a given number of cells. Tumor cells acquire the ability to invade nearby tissue and to disseminate, hence to escape the physiological size limits.
4. **Resource control**. The invasion process is supported by angiogenesis, which is reactivated and maintained to allow the formation of new blood vessels that help to sustain and expand neoplastic growth, and by the ability of adjusting the energy metabolism in order to fuel cell growth and division. As the capacity of the system is limited and the number of cells capable of living in such conditions is bounded, there is a natural “competition” for survival between different cells. The state of equilibrium where each cell has the same possibility to get access to the resources it need for living is lost and the ability to gain an advantage is acquired: the cell might need less nutrients for living by reprogramming the energy metabolism, it can actively harm the neighbours by subtracting nutrients, or it can become capable of exploiting resources that have been recruited by others. Yet, a combination between these “powers” might be advantageous, for instance if two cells both help the other and find resource in them, a mutualistic relationship could be created. All these events are grouped in this functional effect: those that tune how the resources are split among the cells.

We will consider another (non-)functional mechanism for our purposes of modelling:

##### 0. Null effect

It is associated to mutations that do not lead to any observable or functional change in the cell, effectively behaving as neutral mutations.

Different mutations can induce the same functional effect, but possibly with different intensities: for instance, a mutation might lead to the mutation rate being doubled, while another could decuple it and both will be mutations that lie in class 2. Hence a functional capability is defined by two objects: the class of the primary mechanism and its intensity. For this reason we define the *functional event list F* as the set

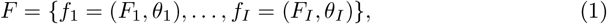

where *I* is the number of functional capabilities included, and *F*_*i*_ corresponds to the class of primary mechanism (one among the 5 we described above) induced by the mutational event, and *θ*_*i*_ indicates the set of parameters, i. e. the intensity. Assuming that each mutation causes an alteration included in *F*, we define a function that associates each genotype *u* = (*u*_1_,..., *u*_*n*_) to its *phenotype*:

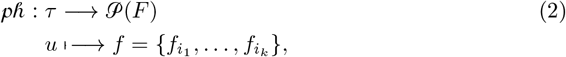

with

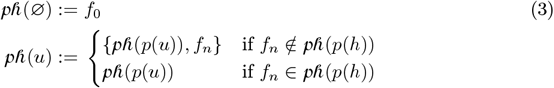

The differentiation among the two cases (1) the newly acquired mutation induces a functional capability already gained through a former mutational event experienced by the parental population -*f*_*n*_ ∈ *ph*(*p*(*h*))- and (2) the newly acquired mutation induces a new functional capability -*f*_*n*_ ∉ *ph*(*p*(*h*))-, is necessary to represent the functional redundancy of the gene network. Some mutations may target different regions of the DNA that encode subunits of the same protein complex. If the complex is already non-functional due to one mutation, a second mutation in another subunit of the same complex does not worsen the defect. Even mutations occurring in different genes that perform similar or overlapping roles within a pathway have a similar behavior: if either gene is mutated, the pathway might be disrupted or overactivated, leading to the same functional effect, while when both genes are mutated, there’s no additional impact because the pathway was already maximally altered by the first mutation. To model additive epistasis, it is enough to duplicate the functional event adding an element to *F* : as said before, the list of functional events *F* is superabundant with respect to the primary characteristics we have described and the reason of that is not only to consider effects of different intensities, but also to better characterize those mechanism that sum up.

The recursive definition of the *phenotype* function given in Eq. (3) sets by default the effect of different mutations as cumulative. In our model, we have chosen to support only additive interactions and neutral epistasis due to the need to balance model complexity with usability as incorporating non-additive forms of epistasis (such as positive, negative, or reciprocal sign epistasis) introduces a significant increase in the number of parameters. This choice is not a limitation of the model’s conceptual framework but rather a deliberate simplification aimed at enhancing its practical applicability. However, our framework remains flexible and extendable: incorporating non-additive interactions is entirely feasible should the need arise or when sufficient knowledge is available to justify the additional complexity.

#### The population evolution model

For each genotype *u* ∈ *τ* we associate *σ*_*u*_ ≥ 0, the birth time of *u*, and (*X*_*u*_(*t*),*t* ≥ *σ*_*u*_) the number of cells, with genotype *u*, at time *t*, the clone. We make the following choices. The process that determines the size of the cell population (*X*_*u*_(*t*),*t* ≥ *σ*_*u*_) is modeled as a size-dependent birth and death process. The theory of size-dependent branching processes, which extends classical branching models by allowing reproduction and survival rates to depend on population size, has its origins in the mid-20th century and counts among its ranks distinguished mathematicians such as Klebaner and Jagers, see [43], [44], [45]. Despite the name, size-dependent branching processes do not satisfy the basic branching property, that corresponds to “Each individual evolves into a branching process independent from and identically distributed to all the others”. Indeed, the law governing a *size-dependent branching process* will depend on the size of the population, which makes it impossible for the evolution of an individual within many, to evolve following the same law as the individual generating the process alone. Recalling the idea that the functional mechanisms of a clone are completely defined by its *phenotype*, the birth and death rates of each population will be phenotype-associated, *a*_*ph*(*u*)_ and *b*_*ph*(*u*)_, i.e. will depend only on the functional effect of the mutations that the clone has cumulated. We can therefore define the process describing the number of cells sharing mutational profile *u* as:

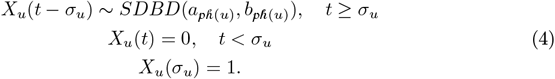

The extension before appearance of population *u* is straightforward as any population will be of zero individuals until its appearance, and then it will start with a single cell, the daughter of a cell with mutational profile *p*(*u*) having acquired an additional mutation. Population *u* arises from cells belonging to *p*(*u*), as the result of a duplication with a new mutational event cumulated in one of the daughter cells. The point process that describes the appearance of the genotype *u* is modeled as a *Doubly Stochastic* point process or *Cox* point process, see [46], with random measure

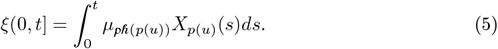

Hence, conditional on *ξ*, the point process is a Poisson process with parameter measure *ξ*. It follows that

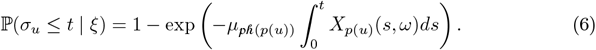

We will set *σ*_Ø_ =0 with probability 1.

The set of genotypes appeared within time t, *N* (*t*)= {*u* ∈ 𝒰 : *σ*_*u*_ ≤ *t*}, is almost surely a Locally Finite Rooted Tree for any *t*, see S1 The model: mathematical details for the proof. Notice that *N* (*s*) ⊆ *N* (*t*) for *s* ≤ *t*.

The stochastic process describing the tumor clonal evolution is

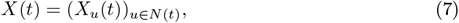

meaning that the state of *X* at time *t* is completely determined once given a LFORT and the vector of the populations abundances, hence *X* : [0, ∞) × Ω → *S*, where the state space *S* is defined by

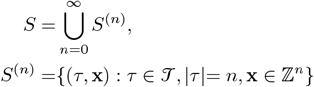

The process *X*(*t*) is a Markov process, see S1 The model: mathematical details for the proof, with generator

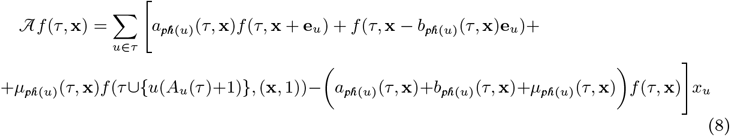

The size dependence of the process is encoded within the of the parameters *a*_*ph*(*u*)_(*τ*, **x**), *b*_*ph*(*u*)_(*τ*, **x**) and *µ*_*ph*(*u*)_(*τ*, **x**) on the state of the process (*τ*, **x**). A comprehensive description of the derivation process of the parameters *a*_*ph*(*u*)_(*τ*, **x**), *b*_*ph*(*u*)_(*τ*, **x**) and *µ*_*ph*(*u*)_(*τ*, **x**) for any given phenotype *ph*(*u*) can be found in S1 The model: mathematical details.

### The simulation algorithm

At each time, the process (*X*_*u*_(*t*))_*u*∈*N*(*t*)_ is a multi-dimensional random variable with dimension equal to the number of populations with at least one alive cell, |*N*(*t*)|. The evolution of each established population follows a size-dependent branching process that can give birth to the first cell of a new emerging population with a new genotype that includes a newly acquired mutational event. The simulation algorithm is a discrete-time approximation of the process (*X*_*u*_(*t*))_*u*∈*N*(*t*)_ which updates all of its components: the abundances at the next time point *X*_*u*_(*t* + Δ) for all *u* ∈ *N* (*t*), the set of established populations at time *t*, and the abundances of the emerging populations, with birth time *σ*_*u*_ between *t* and *t* + Δ. Please note that both the latter updates are needed to correctly refresh the process *X*(*t*), see Eq. (7). The update of the abundances of the established populations at time *t* is computed before the update of the emerging populations as the result of the former is necessary for the calculation of the latter.

A graphical representation of one single simulation step is reported in Fig. S2.1. Moreover, a study for the choice of the largest timestep Δ that controls the approximation error is given, see S2 The simulation algorithm: mathematical details. The pseudocode for the full algorithm is included in S2 The simulation algorithm: mathematical details. In the following, we are describing the main simulation steps. **Established populations update from** time *t* **to time** *t* + Δ. Once the birth parameters have been adjusted using Eq. (S1.11) by substituting the population sizes with the values calculated at the previous time step, *X*_*u*_(*t*) for any genotype *u* emerged by time *t, u* ∈ *N* (*t*), the evolution of population sizes is simulated freezing the birth and death parameters, as if they were constant for the time interval Δ. As there is no ambiguity on the parameters when taking into consideration a single population, we will use *a* and *b* instead of *a*_*ph*(*u*)_ and *b*_*ph*(*u*)_ in this paragraph for the sake of notation simplicity. The distribution of the number of individuals at any time *t* + Δ is well known for classical birth and death processes. In particular, we have that, by temporal homogeneity and setting *P*_*ij*_(Δ) = ℙ (*X*_*u*_(Δ) = *j* | *X*_*u*_(0) = *i*), the following distribution can be derived, see [47]:

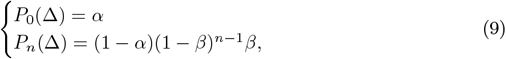

where

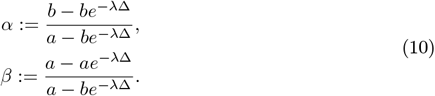

The Embedding theorem, see [48], states that for any choice of discretization step Δ, a birth and death process evaluated at time Δ is a Galton-Watson process with offspring distribution defined by *p*_*n*_ = *P*_*n*_(Δ) given in Eq. (9). This implies that for simulating *X*_*u*_(*t* + Δ), we can employ multinomial sampling with frequencies of occurrences (*p*_1_,..., *p*_*M*_). Since each individual reproduces independently from the others we can consider the evolution of *X*_*u*_(*t*) cells as the replication of *X*_*u*_(*t*) trials of the same experiment whose distribution is given in Eq. (9); the simulation will thus be founded upon:

1. Sample a vector **v** from a multinomial with parameters *X*_*u*_(*t*) and (*P*_0_(Δ), *P*_1_(Δ),…), in order to get **v** (*v*_*m*_ being number of events of type *m*, i.e. birth of *m* individuals, observed after a time interval Δ from a group of *X*_*u*_(*t* + Δ) cells).
2. Update *X*_*u*_(*t* + Δ) = **v** · (0, 1, 2,…).

The simulation error here is driven by the freezing of the birth and death parameters at the beginning of the time step *t*. Taking inspiration from the well-known *τ*-leaping simulation algorithm [49], it is possible to consider a limitation of the time step “ chosen in such a way that the expected state change is small, thus it does not excessively affect the parameters. See S2 The simulation algorithm: mathematical details for a complete study of the simulation error.

#### Birth of new populations

As reported in Eq. (6), the number of daughter populations with parent *u*, born by time *t*, conditional on *ξ* given in Eq. (5), follows a Poisson distribution. This means that the probability of having *k* daughter populations from *u*, in the time interval [*t, t* + Δ) is given by

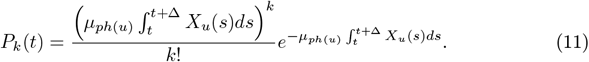

To sample from the correct distribution, it would be required to know *X*_*u*_(*s*) for any *s* ∈ [*t, t* + Δ]. Such information is not available, as we simulate the abundances only at the discrete instants *t* and *t* + Δ. We proceed by approximating the integral with a simple trapezoidal rule. Hence for each population *u*, the number of daughters with a new mutation (i. e. the number of new populations) are sampled from a Poisson distribution with parameter Δ · [*X*_*u*_(*t*)+ *X*_*u*_(*t* + Δ)]*/*2. Each time a new population appears, the new characterizing mutation is named with a unique identifier. The genotype of the new population is then derived concatenating the mother’s genotype with the new unique mutation identifier *v* = *uj*. To each mutational event, a functional capability is associated by sampling a functional event from the functional event list, see Eq. (1), according to an occurrence distribution **r** = (*r*_1_,..., *r*_*I*_). that represents the probability that a random mutational event will result in a given phenotypic effect. For each new population the phenotype is then calculated following Eq. (3)

A user interface developed in `Next.js` allows running the simulator through a configurable environment, where users can set model parameters and execute single simulations. The interface, available together with the source code on `GitHub` (https://github.com/qBioTurin/CancerSimulationInterface.git), returns the full detailed output of a run, including all cell-level information and derived summaries. In Fig. S2.2 and Fig. S2.3 it is possible to see a glimpse of the app.

## Results

The simulator tracks tumor evolution at single-cell resolution, recording for each cell its genotype and phenotype. This information allows for the reconstruction of the complete temporal dynamics and clonal architecture of the synthetic tumor. Several quantitative representations can then be derived at different levels of aggregation.

At the most detailed level, clonal dynamics can be visualized through Muller plots or through evolutionary trees, which depict the temporal expansion of subclones and the mutational relationships among them. Synthetic VCFs are generated as well, by collapsing single-cell genotypes from a random subsample of the population, mimicking bulk sequencing of a biopsy. Each mutation is assigned to distinct reads that are randomly amplified and downsampled to reach a target coverage, optionally drawn from empirical distributions, see Fig. 2. Both the Muller plot, tree plot, and synthetic VCFs are shown in the user interface.

**Fig 2.**
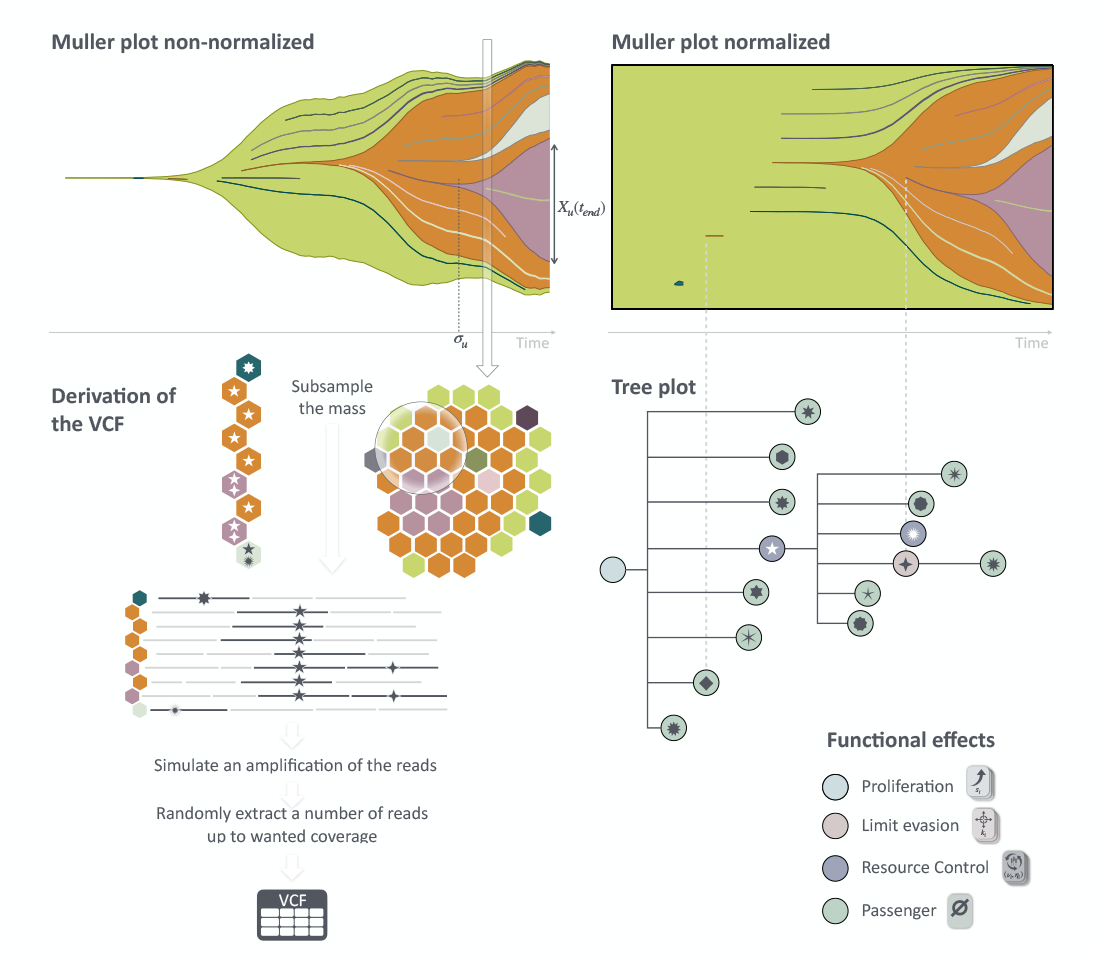
Visualization and re-elaboration of simulated tumors. Muller plots (non-normalized on the left, normalized on the right) provide a comprehensive view of clonal dynamics. Tree plots summarize the evolutionary history, with a color code indicating the functional effects of mutations. In addition, at any chosen time point, the simulation can be re-elaborated to produce a VCF.

At a more abstract level, the evolutionary outcome of each simulation can be summarized by two quantitative indices proposed by [38]: *clonal diversity* 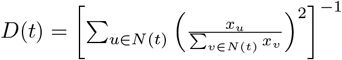, reflecting the average number of clones of the same size, and *clonal nesting* 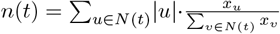, representing the average depth of the clonal tree, which correspond to the average number of mutations per cell. While Muller plots, phylogenies, and VCFs provide detailed insights into individual simulations, the (*n, D*) metrics offer a compact quantitative summary suitable for large-scale comparisons and model validation against empirical data by summarizing the information into a point of a bidimensional plane. Furthermore, these metrics are robust to the addition or removal of rare clones as discussed by the authors, and well established in the literature regarding mathematical modelling for the validation, see [39].

We leveraged both the detailed clonal reconstructions and their (*n, D*) summaries to investigate which evolutionary paradigms naturally emerge from different combinations of functional effects and to validate the model against real data.

### Exploration of evolutionary paradigms

A systematic exploration of the model parameter space has been conducted to characterize the range of evolutionary behaviours generated by different combinations of functional effects and ecological constraints. This analysis, reported in full in S3 Parameter exploration, isolates functional mechanisms and quantifies the resulting clonal architectures across large simulation ensembles. From this broad exploration, a small number of robust and recurring patterns emerge. In the following, we focus on these key insights, showing how the four canonical evolutionary paradigms introduced above arise naturally within the same stochastic framework, without being imposed by construction.

According to *neutral theory* of evolution, the emergence of intratumor heterogeneity does not require a selective advantage, as passenger mutations alone can account for much of the observed variability without conferring any additional fitness to the cell. To test this hypothesis, we ran several groups of simulations, including only passenger functional events. We found that passenger mutations can indeed reach detectable sizes if acquired early enough, although their overall prevalence within the tumor mass remains low. Fig. 3-top-left panel shows an example of clonal evolution restricted to passenger mutations. Most subclones, especially those arising later during tumor development, consist of very few cells. In contrast, a single subclone reached a considerable prevalence. This is not a highly probable outcome (this synthetic tumor is one of the few exhibiting such an event across different simulations), yet it is consistent with the observation that, whenever this occurs, the mutations were acquired very early. These insights are not new. In [23], the authors already showed that, under neutrality, passenger mutations yield a VAF distribution heavily skewed toward very low frequencies, exactly as we observe in our simulations. However, their conclusion that such patterns imply predominantly neutral evolution has been widely criticized [50–52]. Real tumors often display clones with substantially higher prevalence than expected under strict neutrality; the presence of these larger subclones does not necessarily reject the null model presented in [23], but neither proves that evolution is neutral.

**Fig 3.**
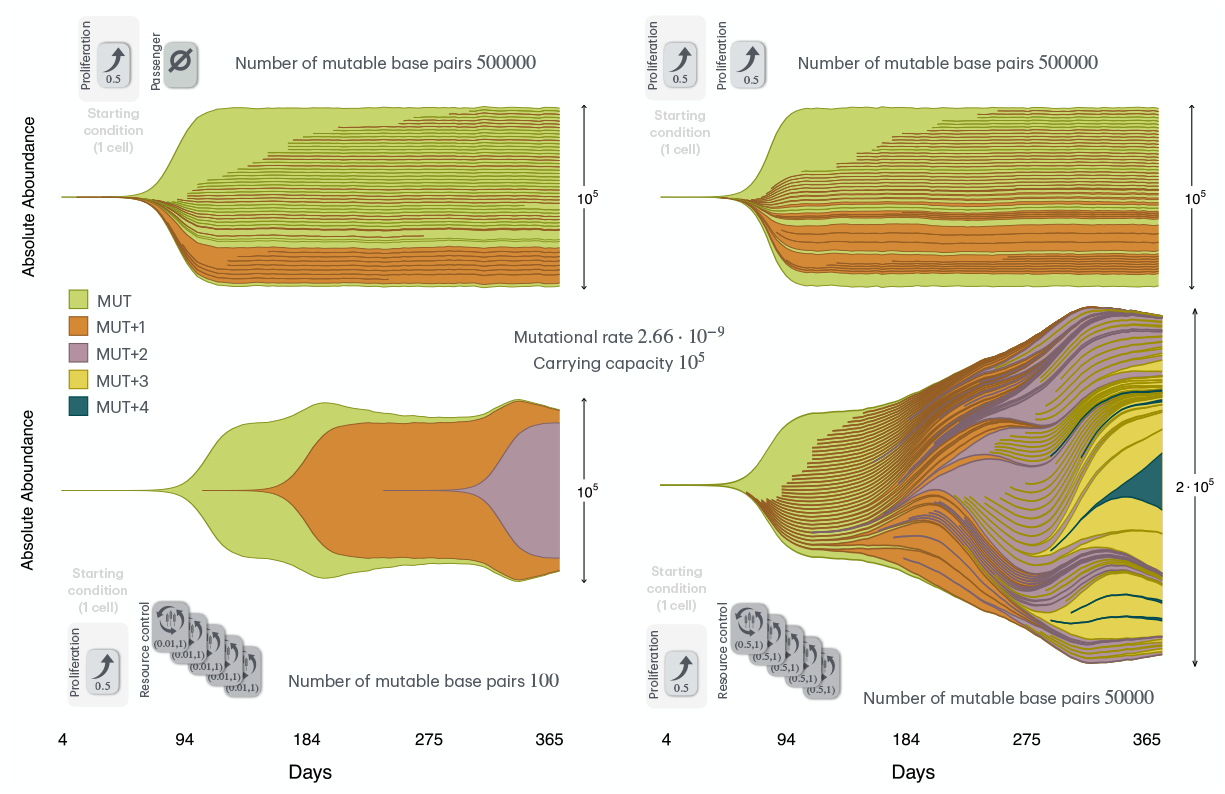
Examples of clonal architectures systematically observed: different evolutionary architectures are captured within our framework and appear to be linked to the functional events. Muller plots are coloured according to the number of mutational events associated with the clone and MUT represent the first oncogenic mutation originating the tumor. The parameters of the model are represented as cards with a symbol corresponding to the type of the primary mechanism of the functional event. When more functional events related to different pathways are considered, one card for each class is represented and the resulting phenotype is calculated according to eq. (3).

What emerges as a novelty, in our model, is that a similar pattern can also be observed for mutations conferring proliferative advantage. As illustrated in Fig. 3-top-right panel, the clonal architectures produced by proliferative mechanisms are strikingly similar to the ones produced by neutral mechanisms, Fig. 3-top-left panel. Subclones appear as thin lines and tend to stabilize at constant relative sizes, once the carrying capacity is reached. This behaviour follows directly from the model rules and can be given a straightforward biological interpretation: when spatial and resource constraints are fully saturated, no clone can expand at the expense of the others unless it acquires the ability either to reduce its own resource demands or to alter the availability of shared resources. There is, however, a clear difference between the two scenarios: the Fig. 3-top-right panel represents a behaviour observed consistently across different simulations: proliferative advantageous mutations appear more likely than passenger mutations to expand up to detectable sizes. These observations show that the type of clonal patterns traditionally associated with *neutral* or *punctuated* evolution can simply emerge when proliferative advantage is the only mechanism allowed and resources are limited: in such conditions, only mutations acquired very early can achieve appreciable prevalence.

By incorporating additional ecological mechanisms of competition and interaction between subclones, the model can generate both *branching* and *linear* evolutionary trajectories, see Fig. 3-bottom panels, without being constrained to any single theoretical scenario. Remarkably, these two considerably different clonal architectures arise from only small changes in model parameters, highlighting the strong sensitivity of the synthetic tumor to the *susceptibility* index. Indeed, both the synthetic tumors shown in Fig. 3-bottom panels are realized by including only functional events of type 4-Resource control, characterized by a neutral (equal to 1) offensive score and a susceptibility index in the range (0, 1). These parameters correspond to a mechanism akin to resistance in which a cell partially escapes the competitive pressure exerted by its neighbours; biologically, such resistance can stem from metabolic reprogramming that enables survival under limited resources or otherwise stressful conditions. When resistance is strong but conferred only by rare mutations, the result tends to be a linear succession of selective sweeps, with each resistant clone replacing all those that came before it. Conversely, when resistance is mild but can be acquired with high probability, multiple lineages persist and expand in parallel, producing a branched architecture.

### Validation against empirical data

To evaluate whether the simulated evolutionary trajectories are biologically plausible, we compared them with real tumor data using the *clonal diversity* (*D*) and *clonal nesting* (*n*) indices evaluated for clones above a minimal frequency threshold of 10^−2^. The idea behind this is that different tumor types occupy distinct regions of the (*n, D*) space, corresponding to characteristic evolutionary behaviours. Real experimental data were obtained from the GitHub repository accompanying [38], where the evolutionary indices of phylogenetic trees previously inferred from multi-region sequencing and single-cell sequencing have been derived. This procedure has been conducted using data from 7 different studies regarding 6 different tumors: acute myeloid leukaemia [53], clear cell renal cell carcinoma [54], mesothelioma [55], breast cancer [56] and [57]), non-small cell lung cancer [58], uveal melanoma and [59]. From each tumor, specific constraints regarding the parameters are derived: for instance, the average lifetime of the cells has been set according to the tissue, they are reported in Fig. 4.

**Fig 4.**
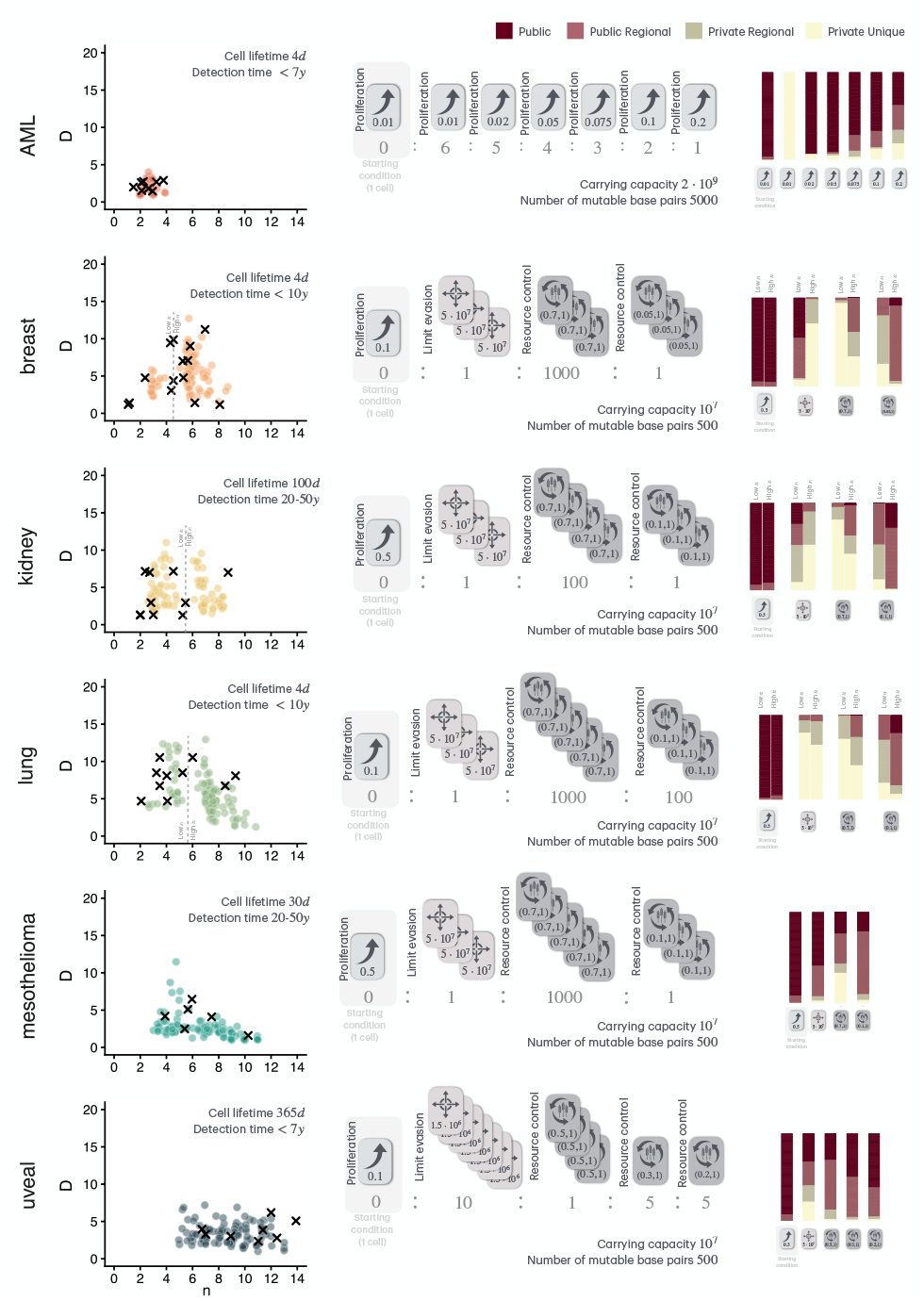
On the **left column**, the x-axis represents the clonal nesting *n* metric and the y-axis represents the clonal diversity *D* metric. Black crosses are the values extracted from real tumors in [38]. Coloured dots are the values extracted from synthetic tumors simulated. The darker and white-bordered dot in each panel is the centroid of the synthetic coloured dots, whose muller plot is depicted beside. On the **central column**, the parameters realizing the best fit according to the likelyhood method, represented as cards with a symbol corresponding to the type of the primary mechanism of the functional event and defining parameters explicitly reported as numbers. The resulting phenotype is calculated according to eq. (3). On the **right column**, barplots describing the distribution of the functional effects according to the prevalence of the mutations sequenced (Procedure described in Fig. 5).

All the simulations have been assumed to start with a single cell carrying a proliferative advantageous mutation (type 1 primary mechanism), that have been excluded by the later acquirable mutations, setting its relative frequency of occurrence *r*_1_ = 0. Passenger mutations (type 0 primary mechanism) are not included in those experiments, as metrics *n* and *D* are defined for driver mutations only. The basic healthy tissue mutation rate per base pair per cell division has been fixed at *m*_*base*_ = 2.66 · 10^−9^ according to [60]. Simulations have been stopped when the synthetic lesion reaches a detectable size, which in literature has been identified with 1 cm^3^ for solid tumors, except for uveal melanoma, which resides in the eye, thus it becomes perceivable already at 1 mm^3^. The size 1 cm^3^ is usually considered to correspond to 10^9^ cells [61]. Though our model only traces alive cells, which are a fraction of the overall tumor mass, hence stopping size has been set to 10^8^ for kidney, mesothelioma, breast, and lung, and to 10^7^ for uveal. For AML, which is a liquid tumor, a larger limit of 10^9^ cells has been used to set the stopping time. Different configurations of primary mechanisms and of their parameters have been tried, running 100 simulations for each chosen configuration.

For each cancer type, a realistic, generous range for the detection time has been set according to literature [62–71], reported in Fig. 4, and different compatible parameter sets have been chosen for the runs. Hence, the (*n, D*) metrics have been calculated for each synthetic tumor (lesion) and each resulting group of 100 points (correponding to the 100 simulations) derived from the same parameter set *θ* has been used to construct a 2-dimensional distribution via kernel estimation 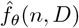. The estimated density 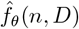 is used to compute the likelihood function of the sampled (*n, D*) true points, 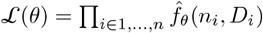. A visual representation of the procedure is depicted in Fig. S3.13. The likelihood value computed is a good estimation of the goodness of fit of the parameter set, hence the parameters realizing the maximum are chosen and represented in Fig. S3.14. In the same Fig. S3.14, for each selected parameter set associated with tumor type, the centroid among all the points (emphasized in the scatterplots) has been extracted and its corresponding synthetic lesion is plotted as a Muller plot and evolutionary tree.

A key distinction emerges between liquid and solid tumors. For AML, the maximum likelihood parameter set consists purely of type 1-Growth-enhancing primary mechanisms, combined with a carrying capacity larger than the stopping size, reflecting the fact that hematologic malignancies can reach detectable sizes without exhausting their environment; here, simple proliferative advantage is sufficient to explain tumor expansion. In contrast, all solid tumors are more likely described by parameter sets that include a carrying capacity smaller than the detection size and acquirable functional events increasing space or resources (type 3-Limit evasion), or exploiting resistance (type 4-Resources control, with susceptibility between 0 and 1 and offensiveness equal to 1). This suggests that the commonly assumed effect of driver mutations as pure growth enhancers may not capture the main mechanisms in solid tumors, where physical constraints and inter-clonal competition play a larger role. Interestingly, once space and resistance are identified as the dominant factors, very different tumor dynamics across solid cancers can be reproduced by only modest changes in parameter values or in the relative frequency of these events. This highlights how subtle shifts in the balance of space acquisition and resistance can generate diverse clonal architectures, reflecting the heterogeneity observed across different tissue types and environmental contexts.

Together, these results validate the simulator as a faithful model of tumor evolution. Its fitted configurations recover the qualitative differences between liquid and solid cancers and reproduce realistic tumor sizes and growth times. Moreover, the fit to empirical data reinforces the conclusion drawn from the sensitivity analysis: in solid tumors, evolutionary dynamics are primarily shaped by resource limitations and by the ability of subclones to evade or withstand ecological constraints, rather than by proliferative advantage alone.

### Synthetic multiregional sequencing from multiple tumors

Following [24], we classified mutations according to their spatial pattern of occurrence and analyzed these patterns in relation to the associated functional events.

For each cancer type, we considered the parameter set maximizing the likelihood of the 100 simulated tumors. For each tumor, we mimicked a multiregional sequencing following the procedure described in Fig. 2, repeated 10 times over non-overlapping regions of 10^4^ cells. Relying on the individual mutation IDs, mutations detected in all (or all but one) sequenced regions were classified as *Public*; mutations detected in a single sequenced region as *Private Unique*; and mutations shared by some but not all sequenced regions as *Regional*, further subdivided into *Public Regional* (present in 4 − 8 regions) and *Private Regional* (present in 2 − 3 regions). For each functional event, we counted the number of mutations belonging to each class (Public, Public Regional, Private Regional, Private Unique). The resulting contingency tables from the 100 tumors of the same kind and sharing the same parameters (and thus functional events) were then aggregated into a single frequency distribution, which shows how different kinds of functional capabilities are associated to distinct spatial patterns inferred from the simulated VCF data. The multiregional sequencing procedure is described in Fig. 5, while the results are shown in Fig. 4. For breast, kidney, and lung cancers — where simulations plotted on the (*n, D*) plane revealed a separation along the horizontal axis - we preformed the analysis separately for low-*n* and high-*n* synthetic tumors and compared the resulting distributions.

**Fig 5.**
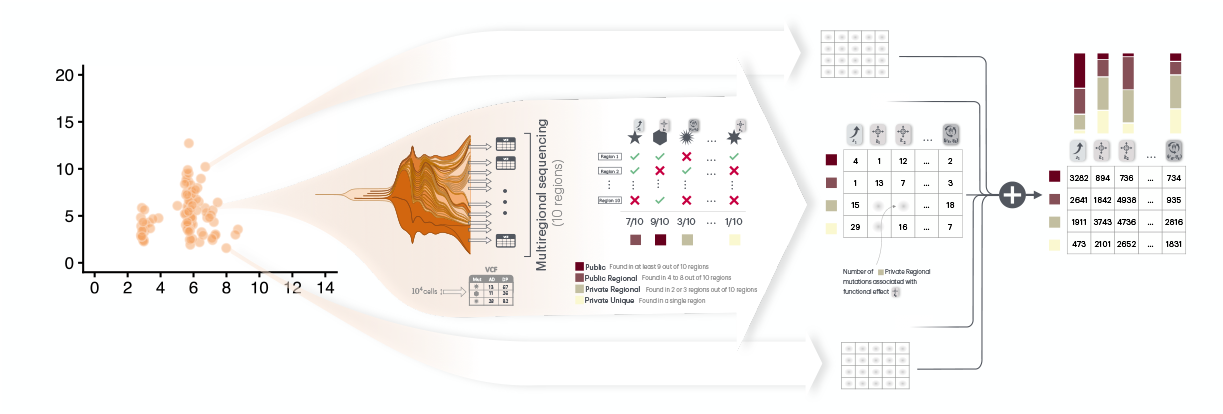
Procedure of multiregional sequencing used for the classification of the mutations based on spatial pattern of occurrence and analysis of these patterns in relation to the associated functional events.

AML evolution is solely driven by mutations or structural variations that directly affect proliferative capacity. Interestingly, in breast and kidney cancers, the distribution of mutation classes differs depending on whether tumors are characterized by a low-*n* or high-*n* number of driver gene mutations. Specifically, the last column of Fig. 4 reports a snapshot of the distribution of mutation classes (Public, Public Regional, Private Regional, Private Unique), grouped by functional event, at the time when the tumor reaches a detectable size (diagnosis time). We see two evolutionary scenarios. In the first scenario, tumors with a low-*n* number of driver mutations are characterized by public mutations associated with type 3-Limit evasion functional effects. Moreover, we could assume that those mutations have been acquired during the early phases of cancer progression, see Fig. S3.15 where the times to diagnosis for the two groups are plotted and the low-*n* group is shown to have significantly smaller times to diagnosis. Notably, the three distinct categories of functional effects can be related to different biological processes involving genes linked to pathways associated with loss of cell adhesion and/or increased efficiency of nutrient usage and survival under hypoxia or starvation. Distinctly, the tumors characterized by a high-*n* number of driver mutations exhibit a different evolutionary pattern. These tumors show a higher prevalence of private mutations related to type 3-Limit evasion functional events, alongside a more persistent presence of public mutations associated with type 4-Resource control functional events. As described by Fig. S3.15, the high-*n* tumors are associated to larger times to diagnosis, hence to slower growth trends. This suggests that, from a biological perspective, alterations that induce type 4-Resource control functional capabilities may be acquired more slowly over tumor evolution. For these tumors, seven (kidney) and six (breast) distinct functional events within the type 4-Resource-control category are modeled, characterized by different degrees of susceptibility and offensiveness. These events can be interpreted as related to internal resource generation during starvation or stress, support of rapid proliferation and membrane remodeling, or reduced immune metabolic competition, ultimately leading to increased resource availability for cancer cells.

A similar analysis was performed for lung cancer, stratifying tumors into groups with low-*n* and high-*n* number of driver mutations. In this case, intratumoral heterogeneity appears to be more pronounced, as more cases result with large *D* values, see Fig. 4. The type 3-Limit evasion functional events are mainly Private unique classified, both for the high-*n* and low-*n* groups. This suggests a more complex and heterogeneous evolutionary landscape in lung cancer, where multiple functional strategies may coexist at the time of diagnosis.

Finally, in mesothelioma and uveal melanoma, the number of driver mutations is more concentrated within a narrower range and is associated with limited heterogeneity, as reflected by lower values of the heterogeneity parameter, *D*. These tumors are characterized by a generally high proportion of public mutations, broadly distributed across all the biological categories considered, indicating a more homogeneous evolutionary trajectory dominated by shared functional events.

## Discussions

We have designed and implemented a stochastic tumor evolution model that unifies, within a single mathematical framework, the main conceptual approaches currently proposed in the literature. In particular, our model integrates branching process descriptions of cell division and mutational events [12,15, 33–36], neutral expansion scenarios as formalized in the Big Bang model [23, 24], and ecological perspectives that emphasize environmental limitations and clonal competition [12, 13, 37–39]. By explicitly incorporating resource limitation and density-dependent feedbacks, our framework extends these models beyond unconstrained growth while retaining a fully stochastic core; moreover, it allows the acquisition of mutations that provide different kind of advantages, in line with recent theories of cancer hallmarks [40–42].

Using our model, we performed a large set of systematic simulation experiments to quantify how the evolutionary fate of newly arising subclones depends on both the functional nature of the mutation and the phase of tumor growth at which it is acquired. Specifically, we compared the survival probability and final prevalence of subclones carrying neutral mutations to those acquiring functionally disruptive mutations of different classes (i.e. proliferative advantage, evasion of growth limits, and control or modulation of shared resources) stratifying results by early expanding versus late saturated growth phases of the mass.

These experiments consistently revealed a biphasic pattern of tumor evolution. In an initial phase, before the expanding mass reaches the physical and metabolic limits of the surrounding tissue, proliferative advantages can transiently influence clonal dynamics, although they remain confined to low-prevalence ranges. In the subsequent phase, once the available space and nutrients are fully exploited, only mutations that alleviate environmental constraints, such as resistance to resource scarcity or the ability to bypass local limitations, allow clones to escape saturation, continue expanding, and potentially dominate the population.

Previous stochastic models that aimed to describe tumor growth using a branching-process approach centered exclusively on driver mutations that confer proliferative advantage, see [12,15,33–36]. More recent frameworks, such as those proposed in [38] and [39], have adopted an explicitly ecological perspective, interpreting environmental limitation primarily as spatial confinement. Their models elegantly show how local crowding and limited dispersal can influence clonal competition, providing a convincing explanation for the structure of tumors growing in glandular or compartmentalized tissues. Our approach shares this ecological foundation but diverges conceptually: we argue that it is not the geometry of space itself that governs tumor evolution, but rather the control and distribution of shared resources. In our framework, competition arises from differential access to these resources and can be modulated by new driver mutations, thus the creation of new spatial compartments is not required to explain clonal diversity. Still, these spatially explicit models remain valuable, and for tumor types where physical architecture plays a dominant role, their principles could be seamlessly integrated into our simulator, further demonstrating its flexibility and generality.

In conclusion, the proposed framework provides a flexible and integrative approach to exploring tumor evolution by jointly analyzing heterogeneous data sources routinely generated in both clinical practice and research settings. These include longitudinal clinical monitoring data, targeted mutational panels, and whole-exome or whole-genome sequencing data. By integrating these data streams, the framework enables a systematic assessment of how empirical observations intersect with the results of simulation-based evolutionary models. At the same time, the framework allows domain-specific knowledge for a given cancer type to be incorporated into the analysis, supporting hypothesis-driven exploration of tumor evolution. Through an intuitive graphical interface, users can visually inspect evolutionary trajectories and identify the classes of mutational effects that drive distinct tumor progression patterns. This combination of data integration, simulation, and interactive visualization enhances interpretability and provides a practical tool for linking molecular alterations to evolutionary dynamics in cancer.

---

**Figure suppl S1_1.**
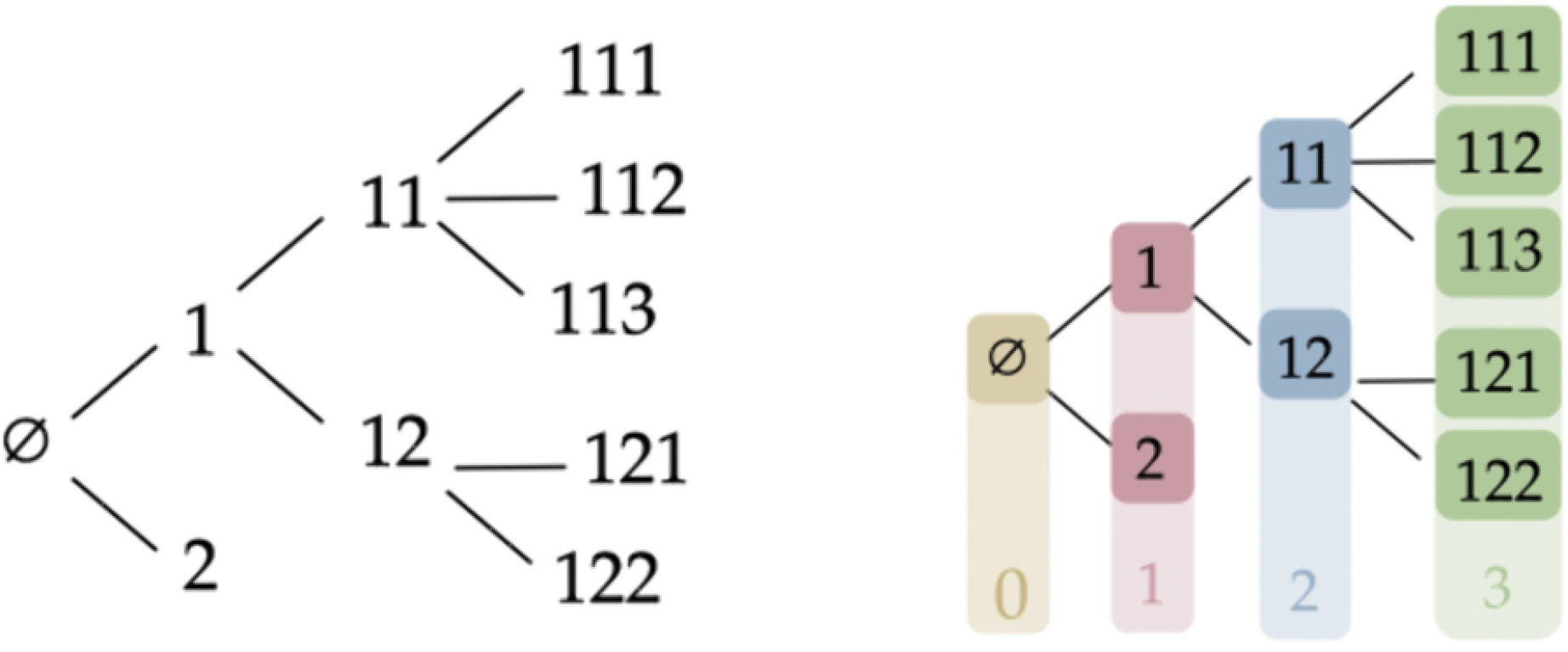

**Figure suppl S1_10.**
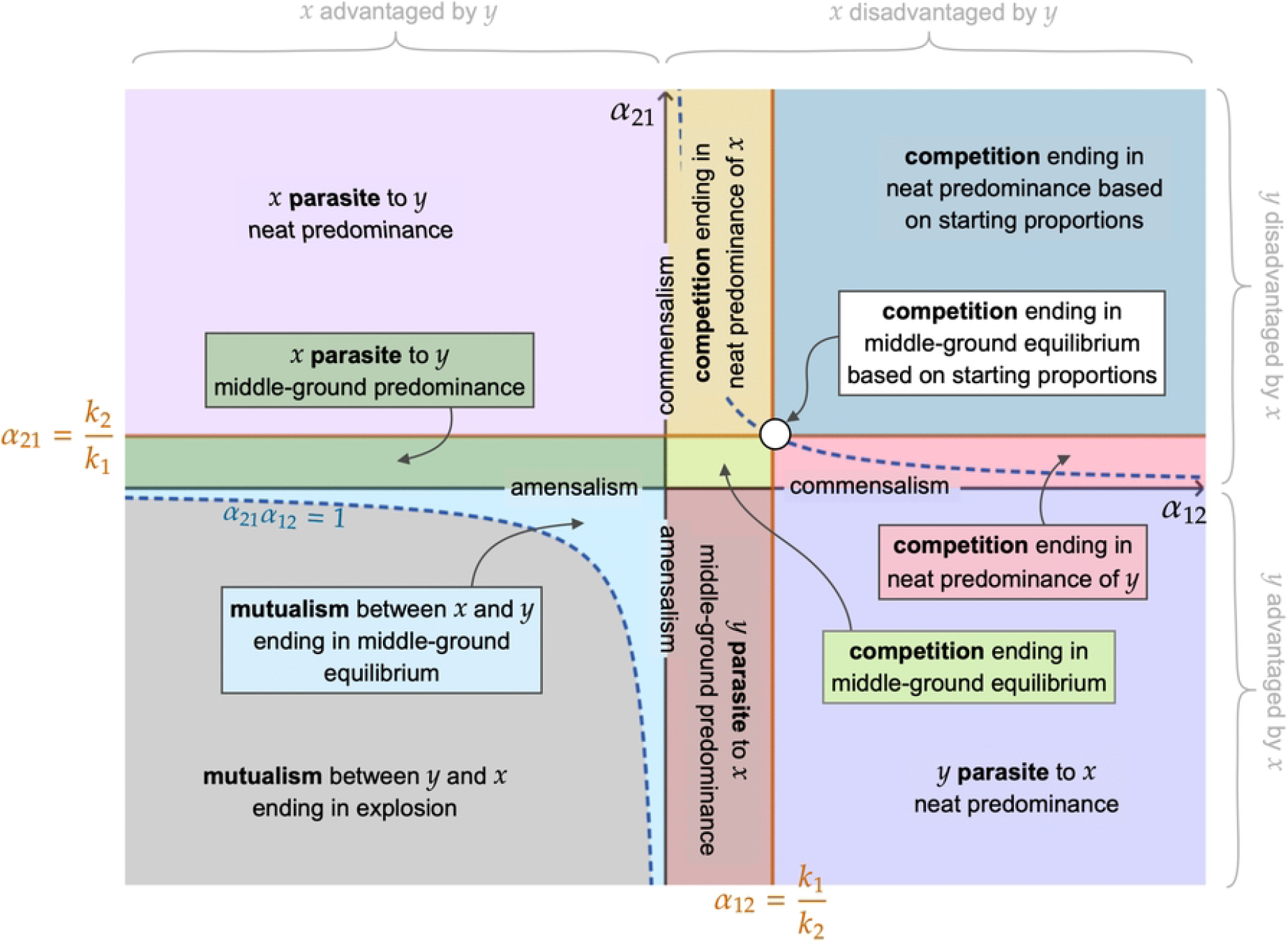

**Figure suppl S1_11.**
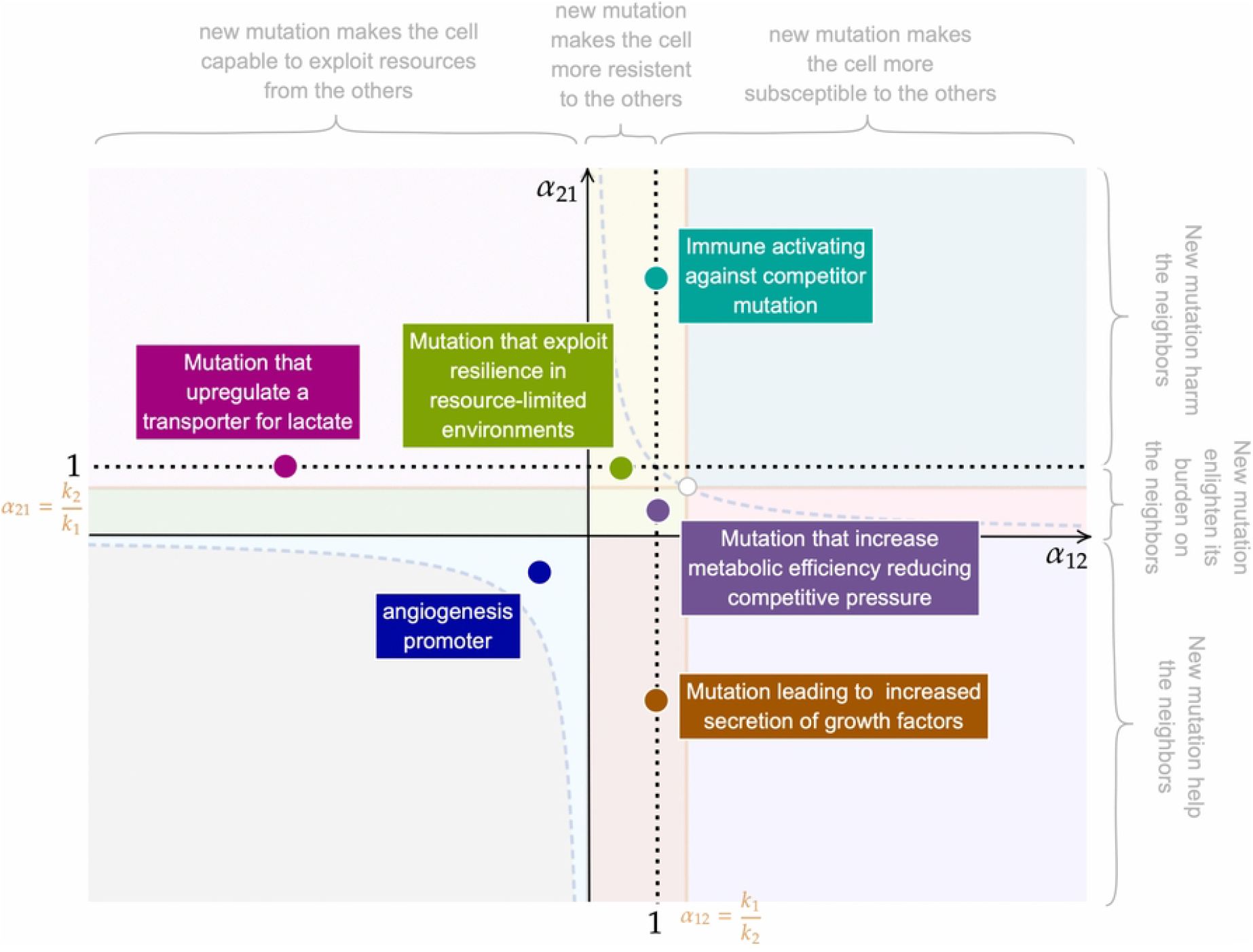

**Figure suppl S1_2.**
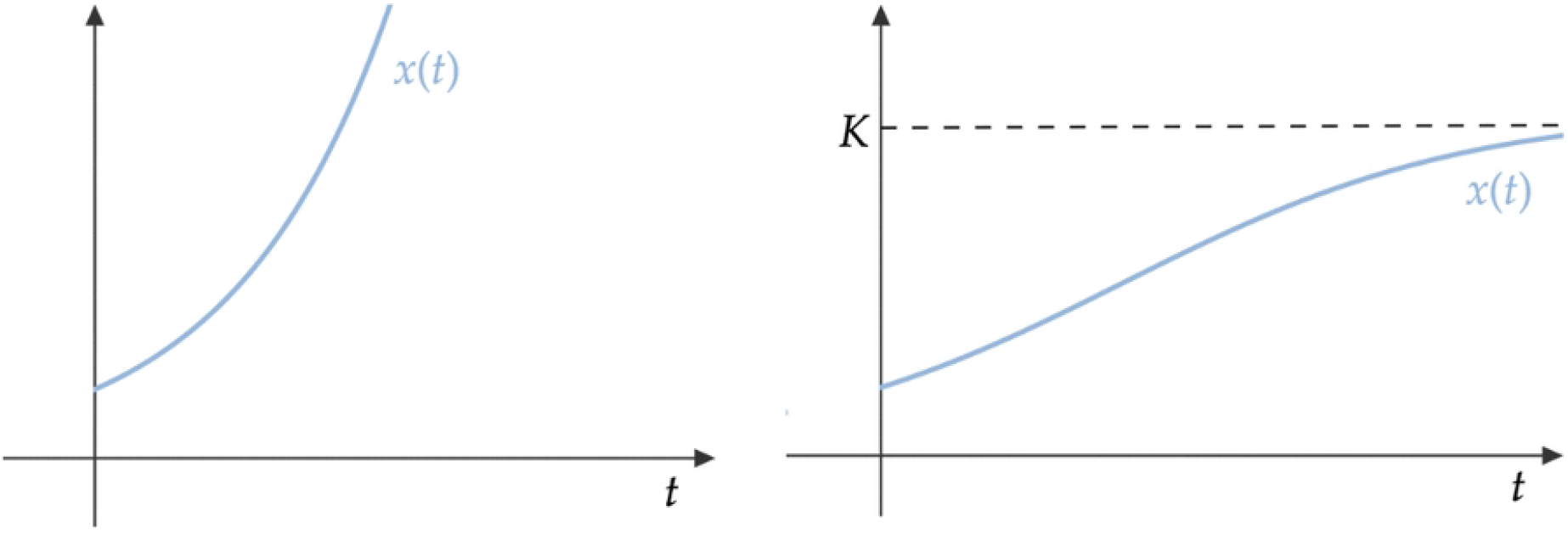

**Figure suppl S1_3.**
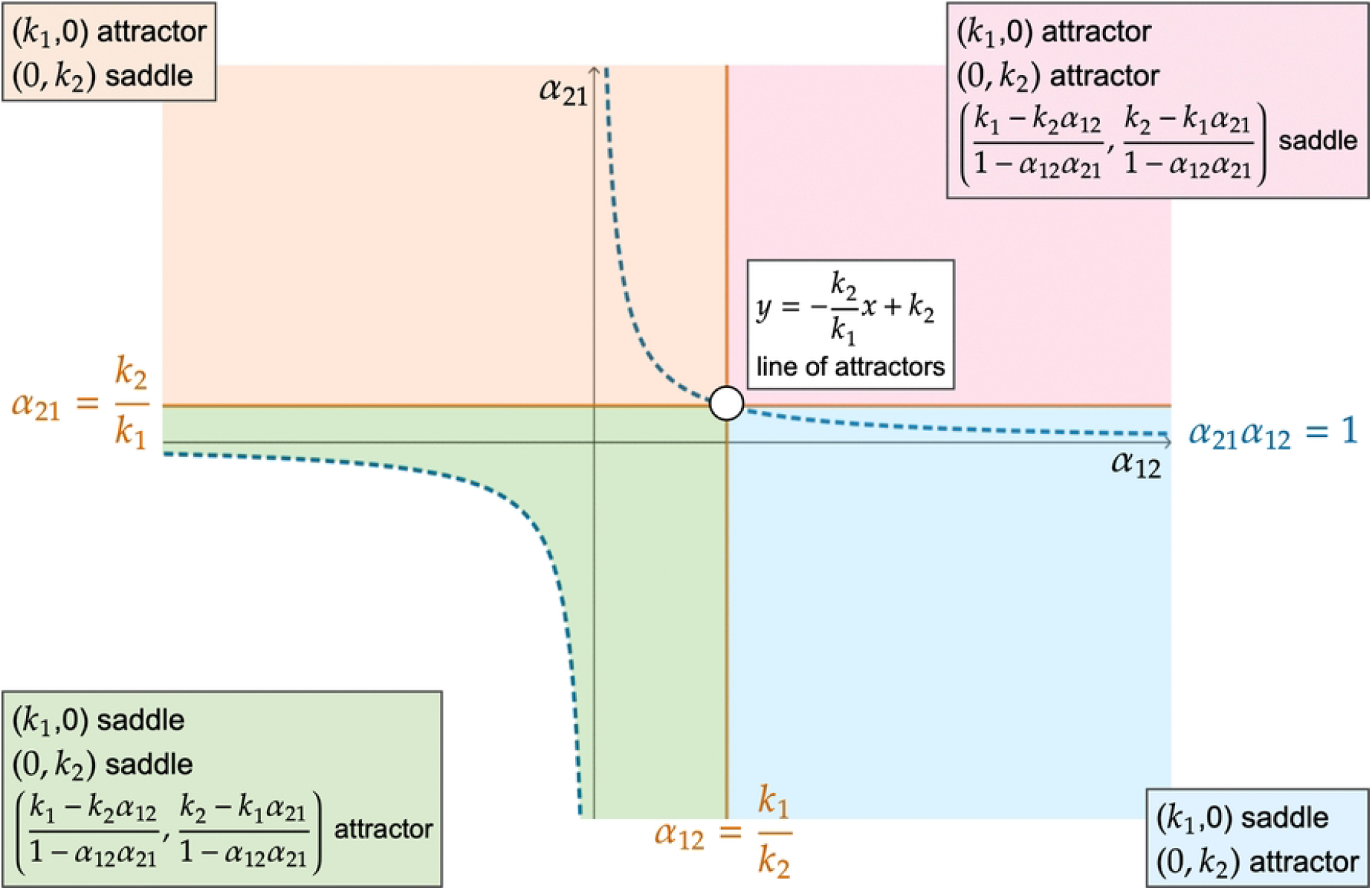

**Figure suppl S1_4.**
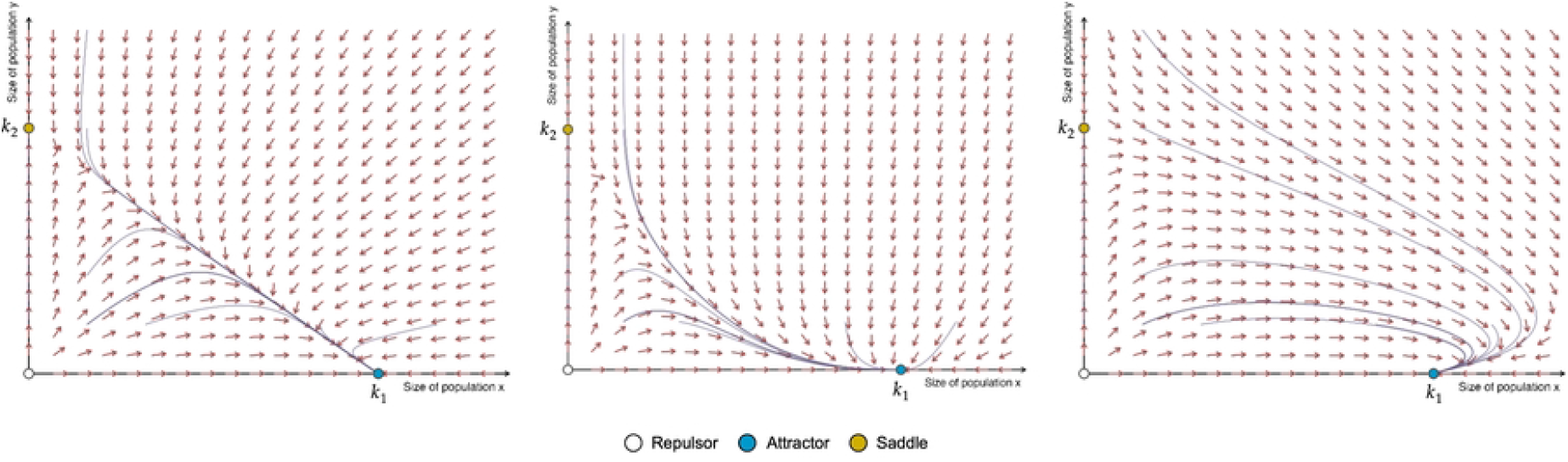

**Figure suppl S1_5.**
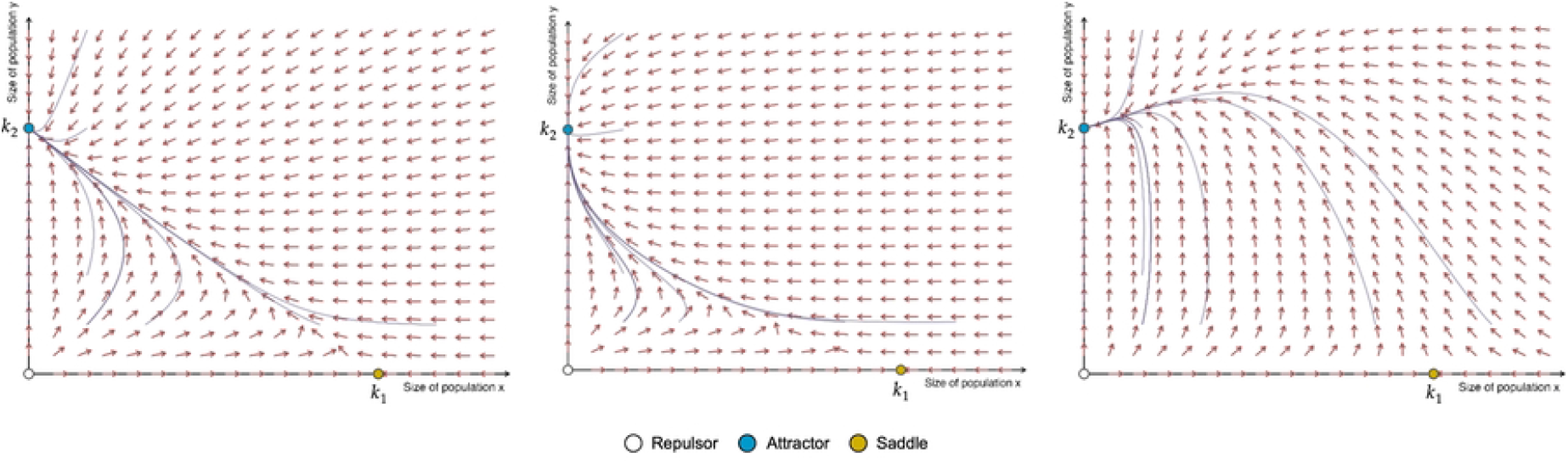

**Figure suppl S1_6.**
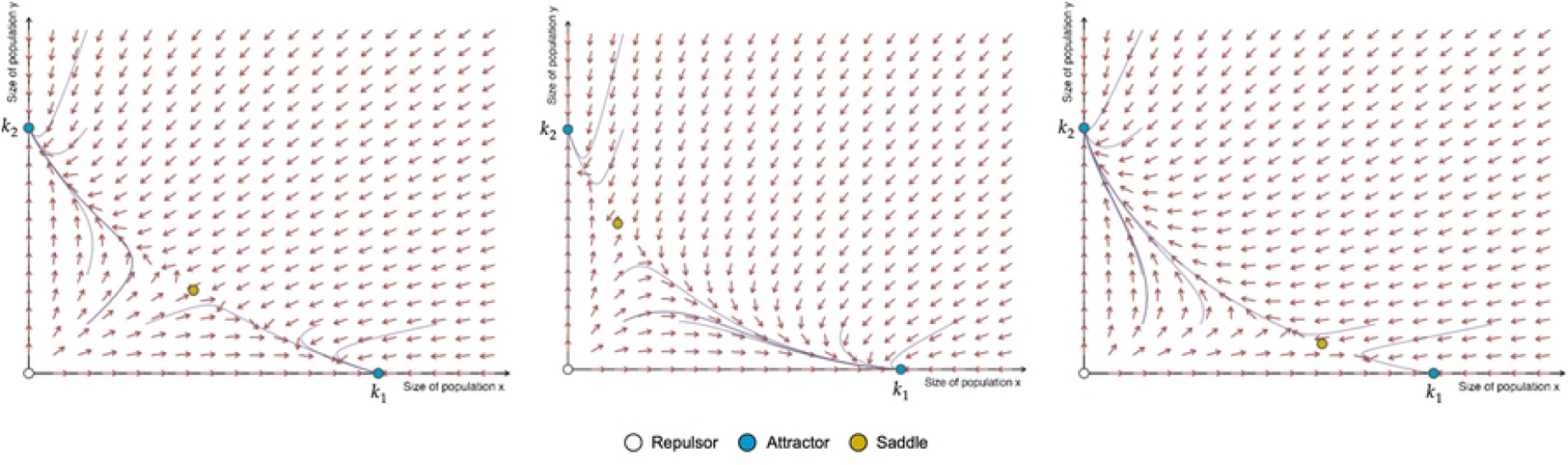

**Figure suppl S1_7.**
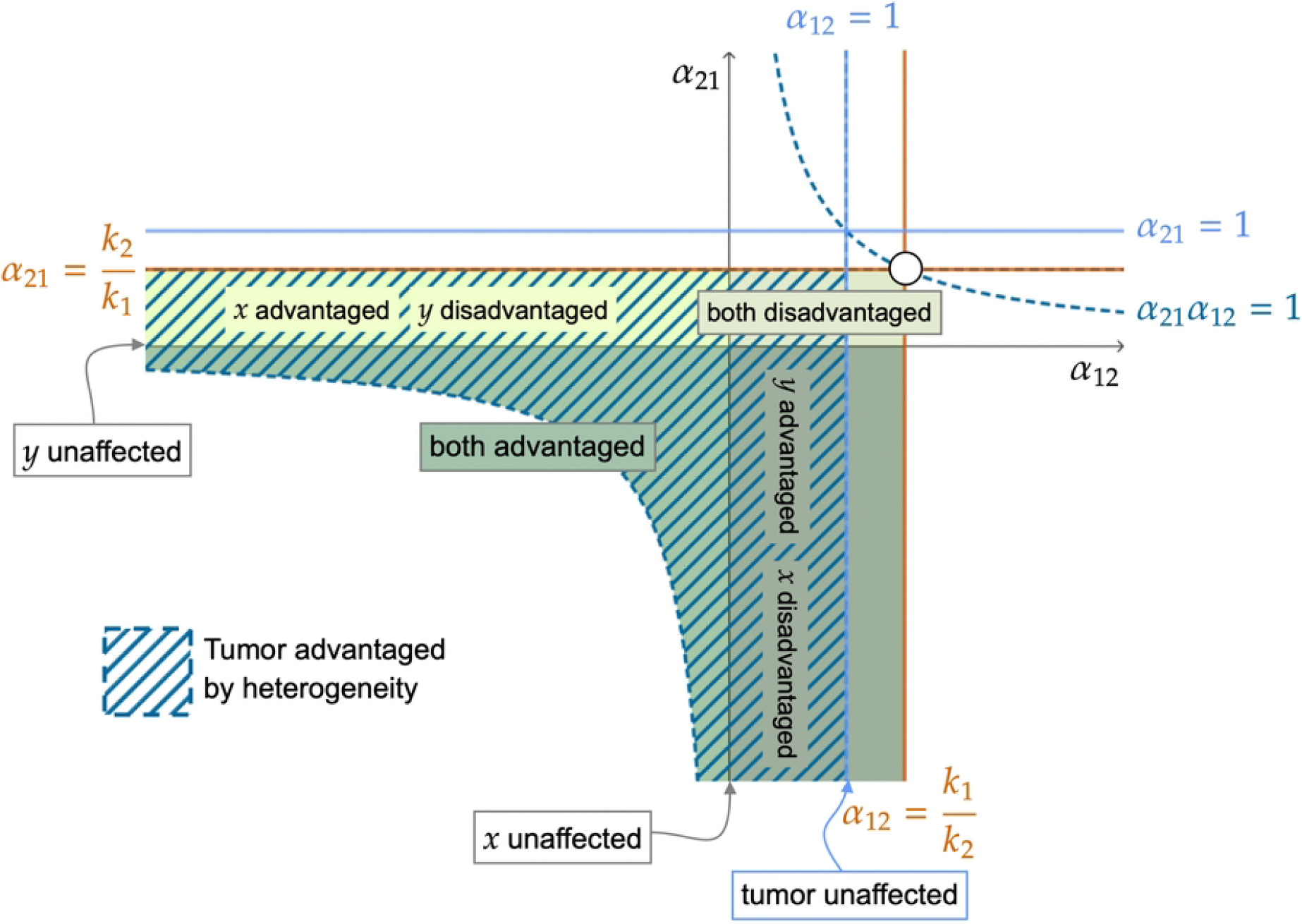

**Figure suppl S1_8.**
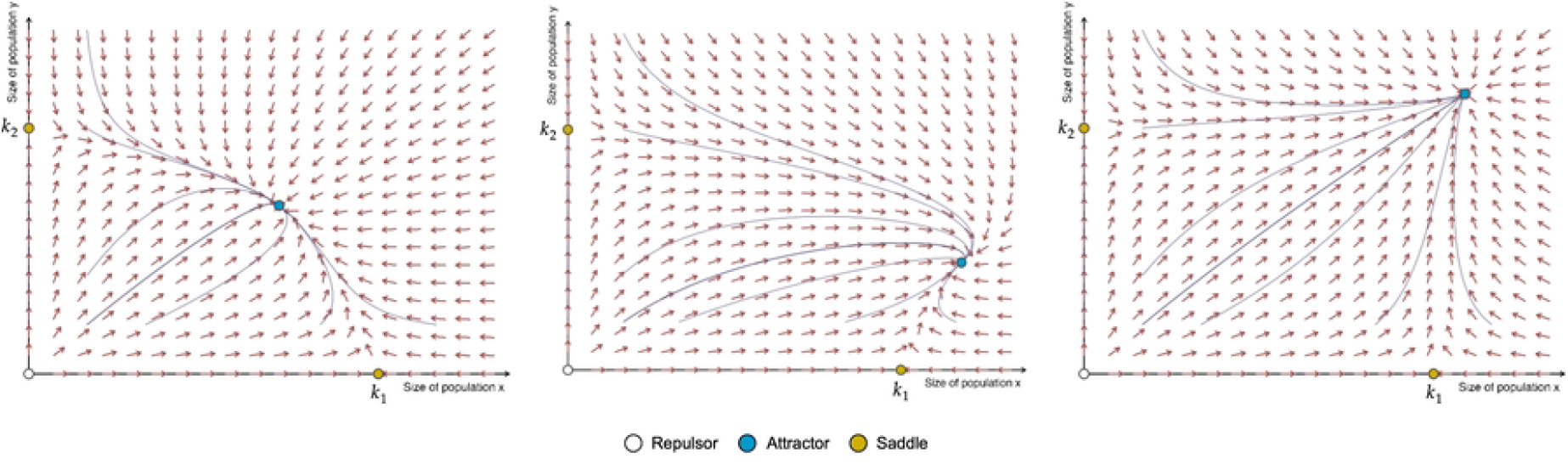

**Figure suppl S1_9.**
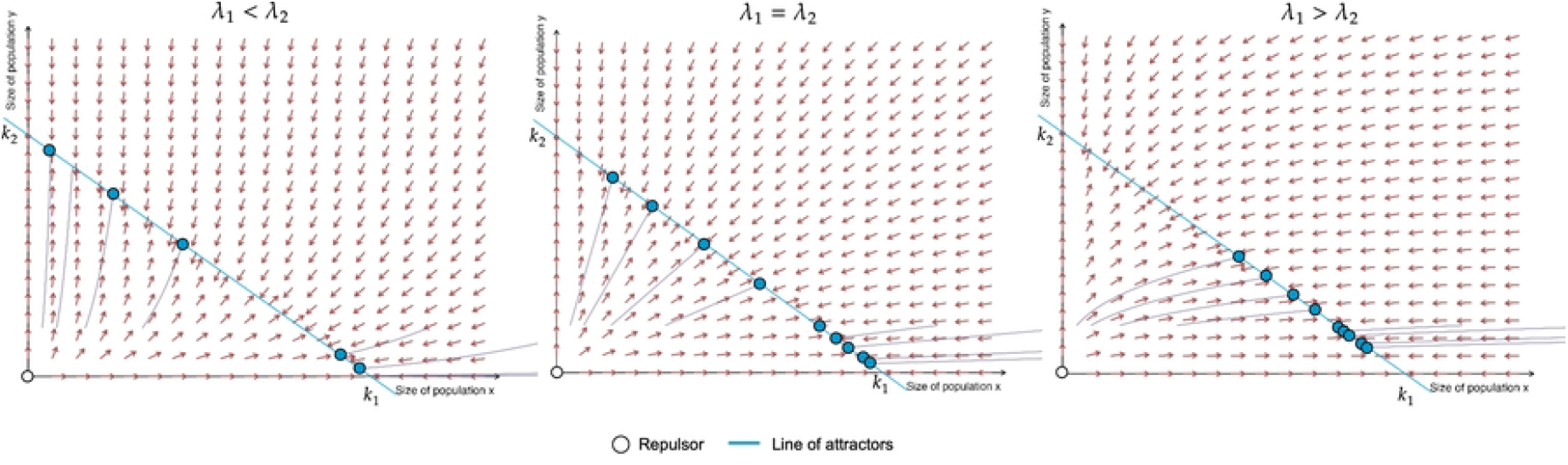

**Figure suppl S2_1.**
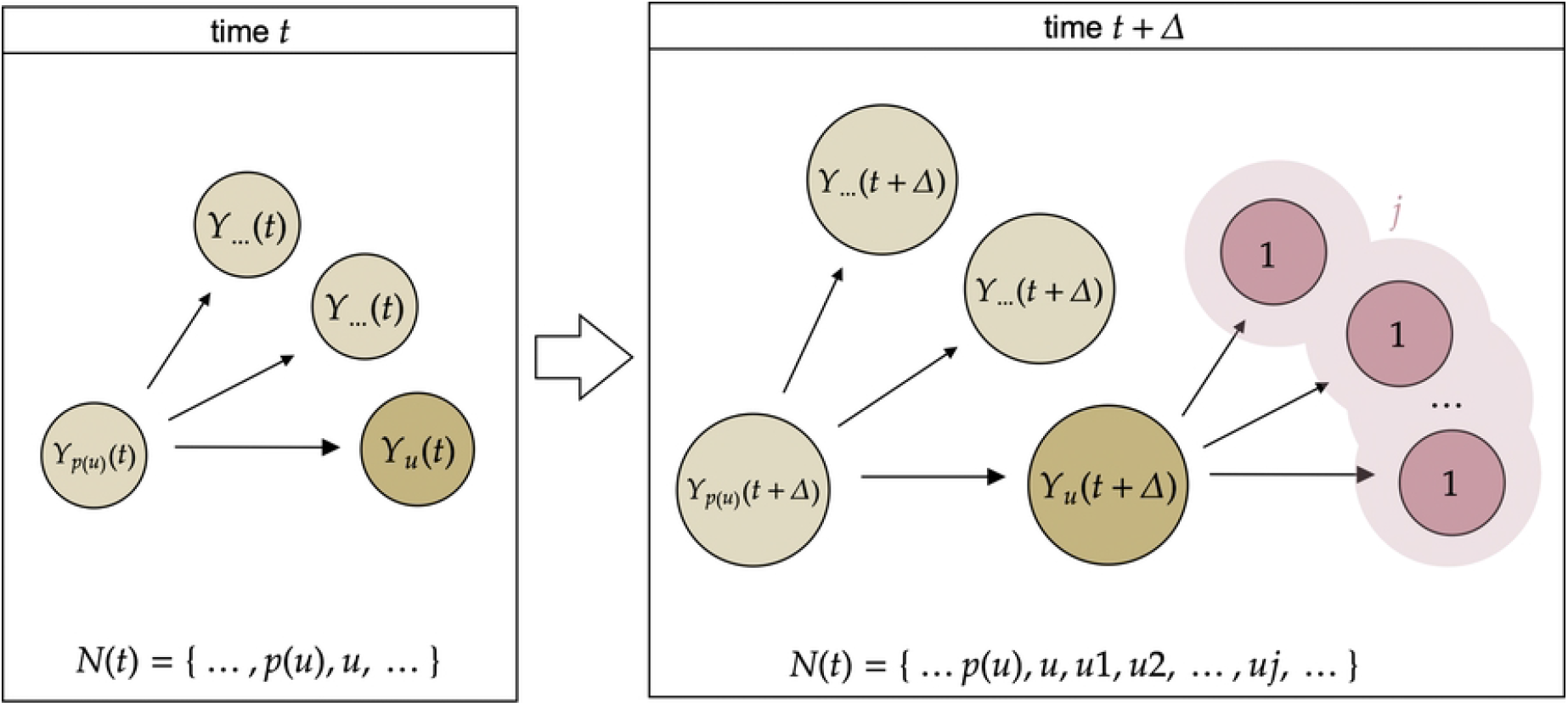

**Figure suppl S2_2.**
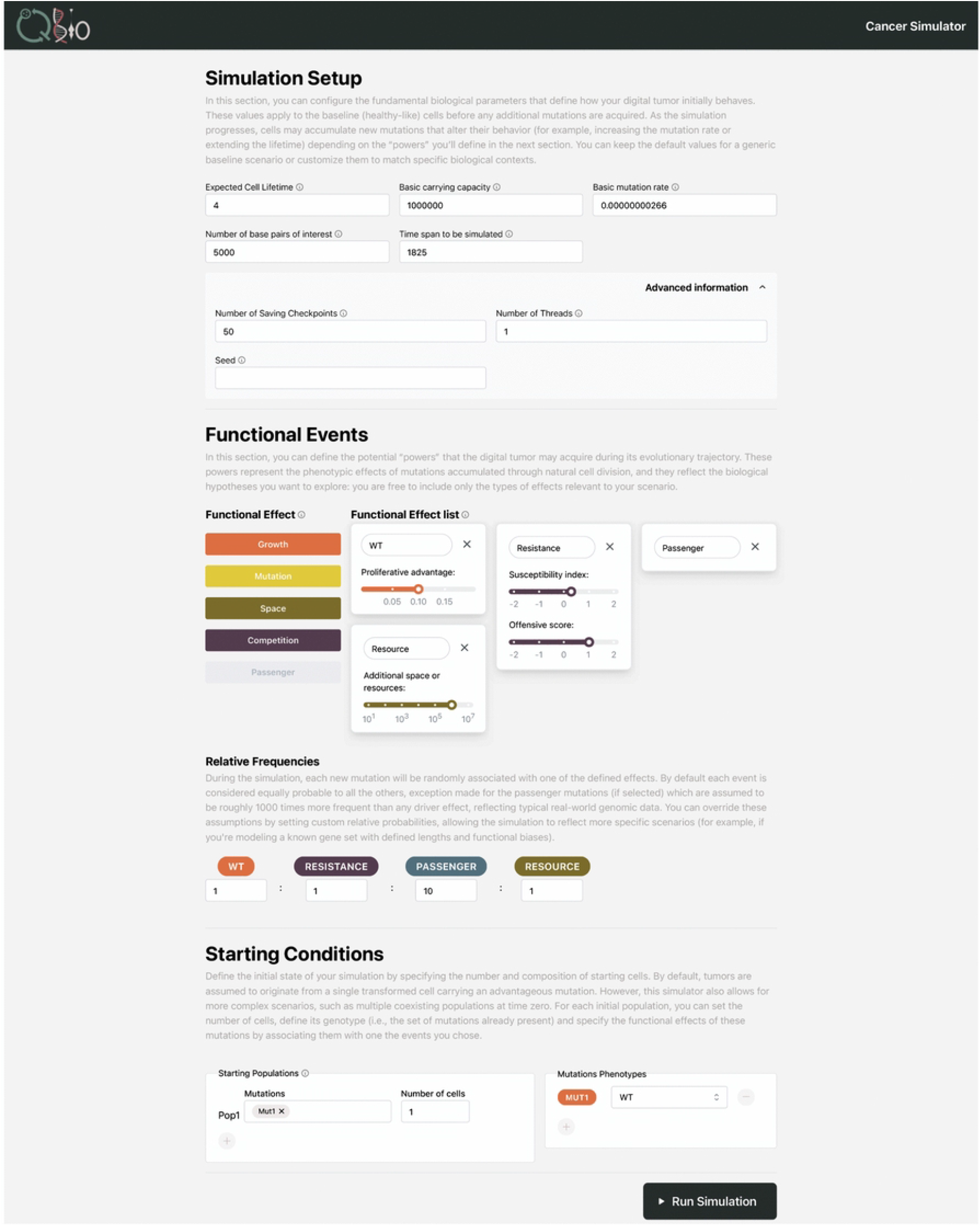

**Figure suppl S2_3.**
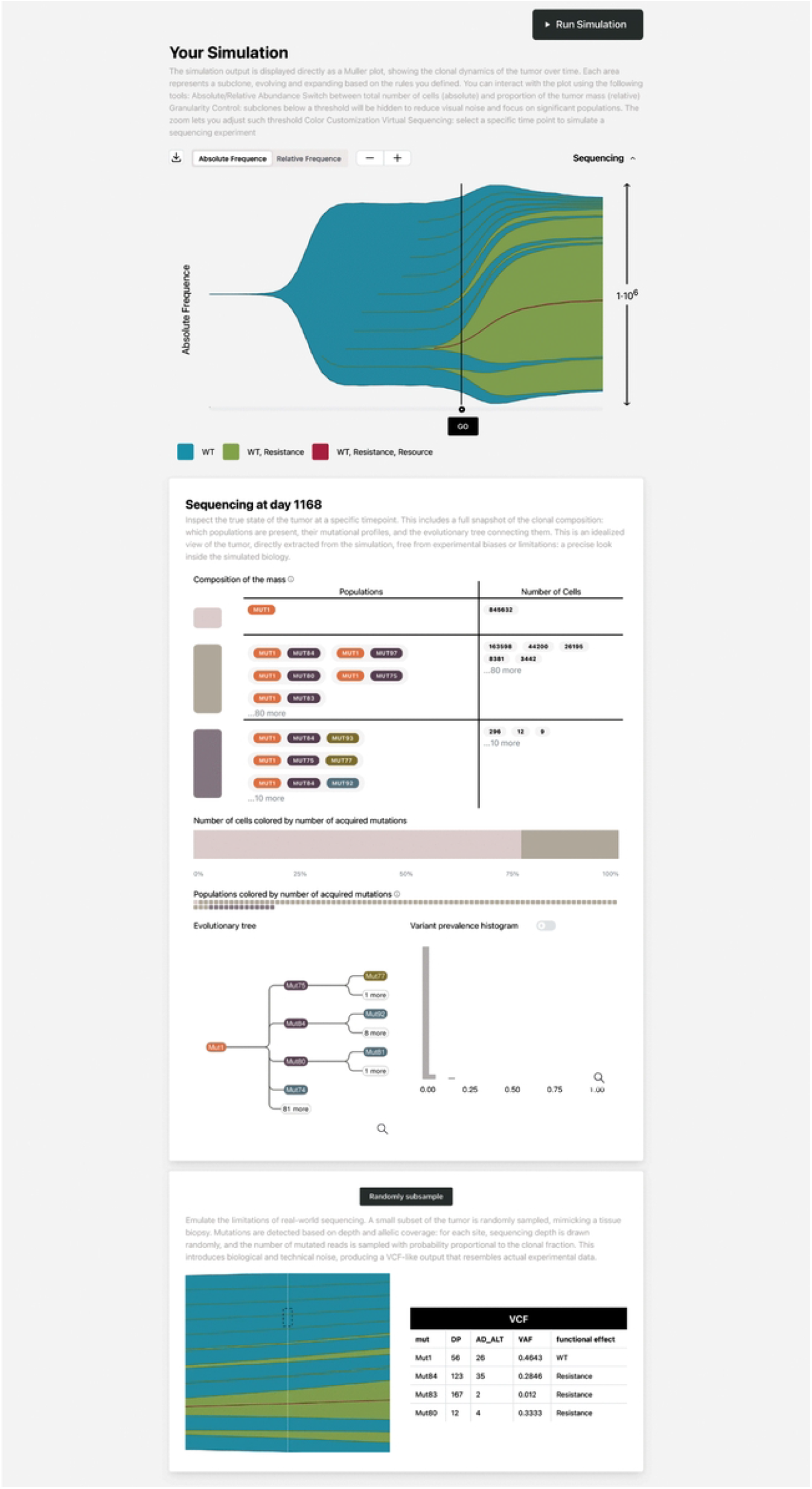

**Figure suppl S3_1.**
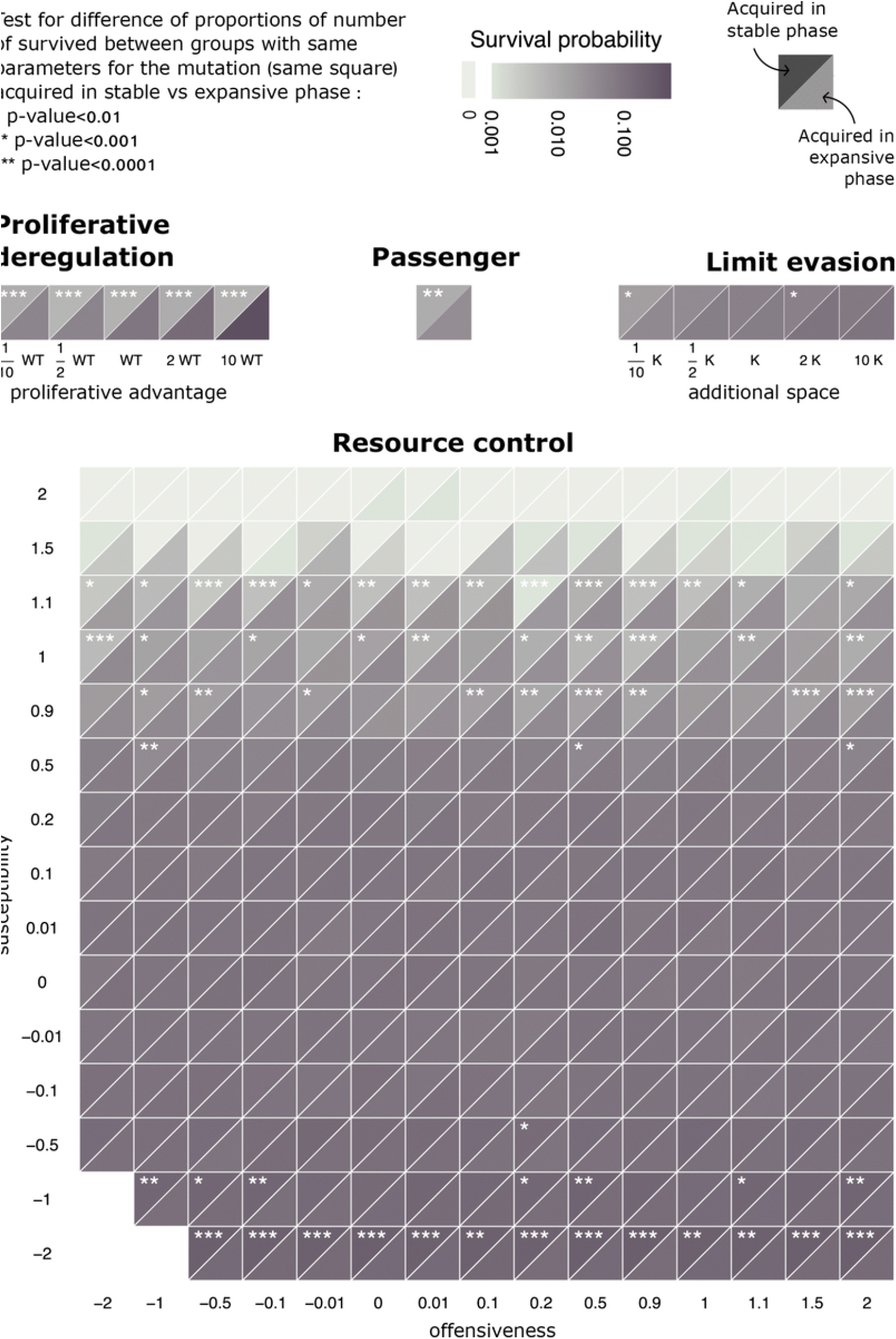

**Figure suppl S3_10.**
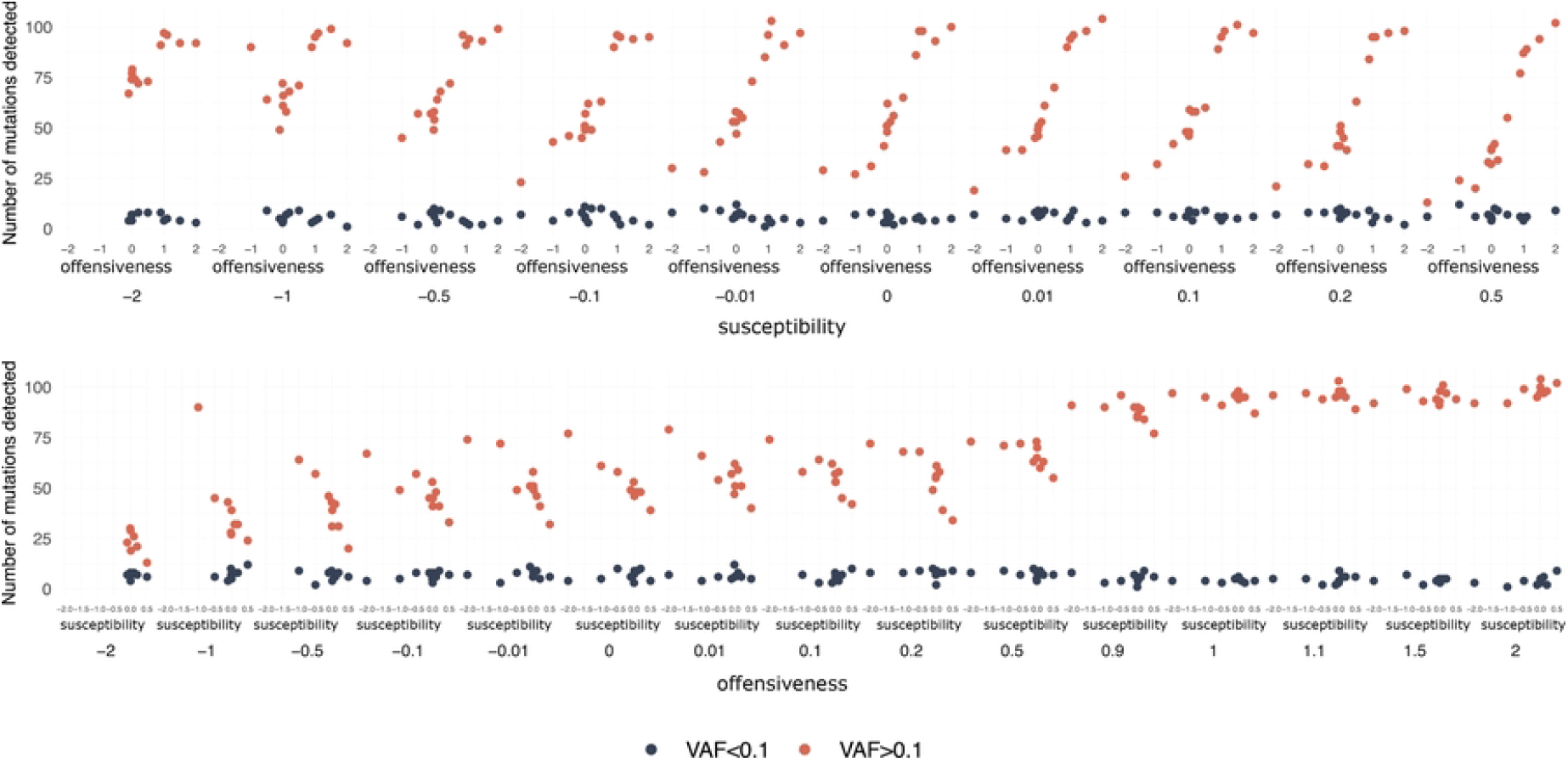

**Figure suppl S3_11.**
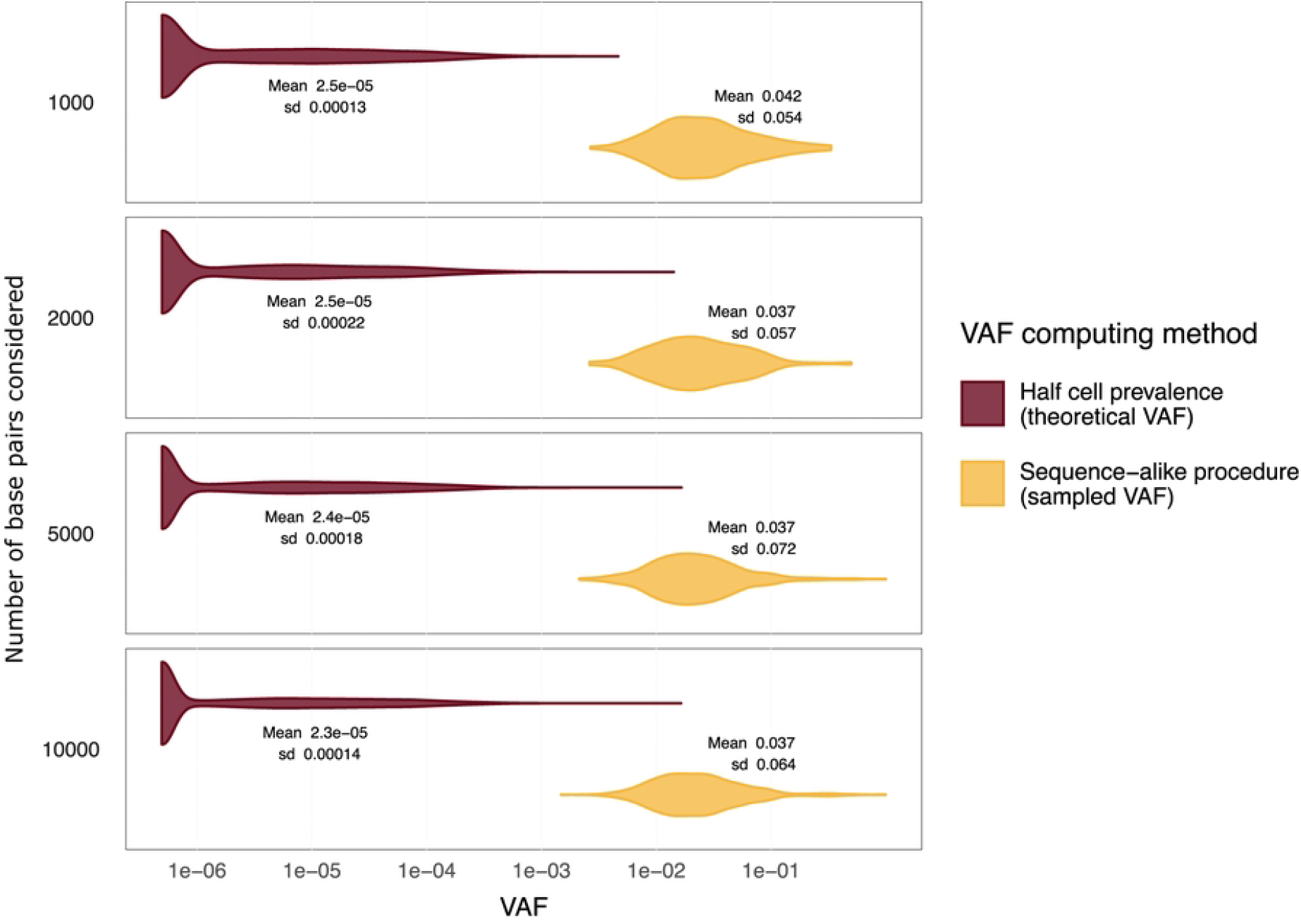

**Figure Figure suppl S3_12.**
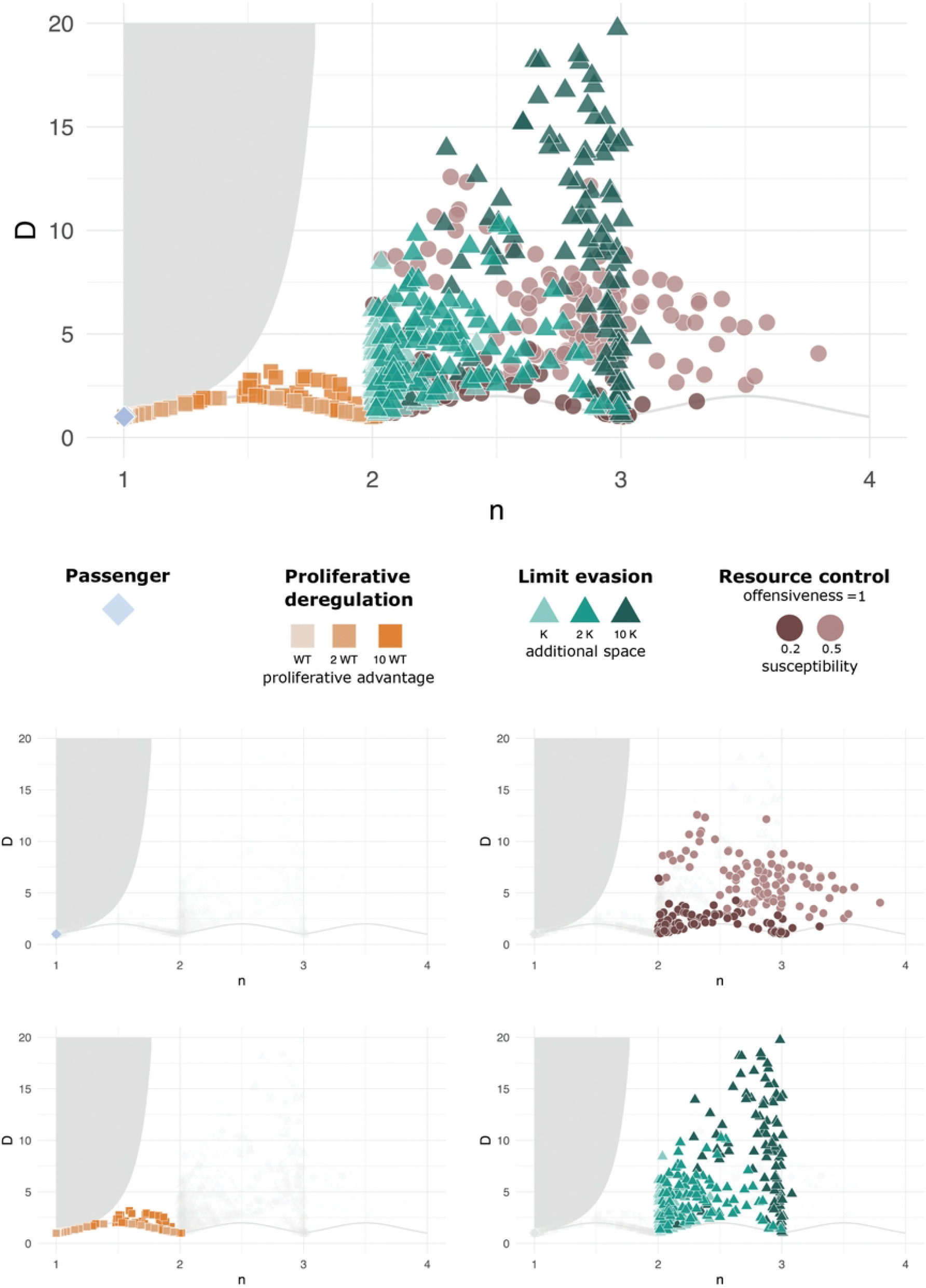

**Figure suppl S3_13.**
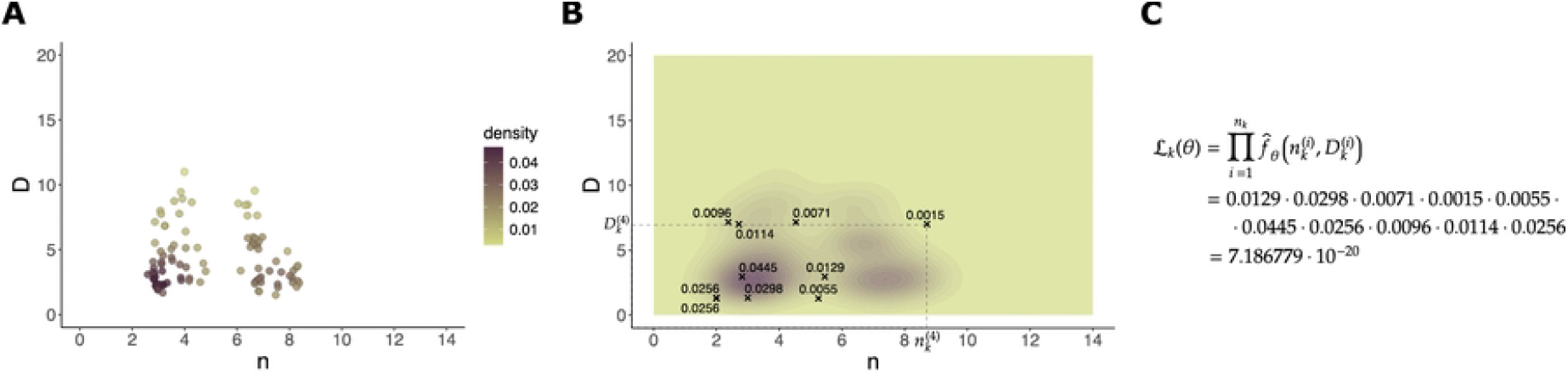

**Figure suppl S3_14.**
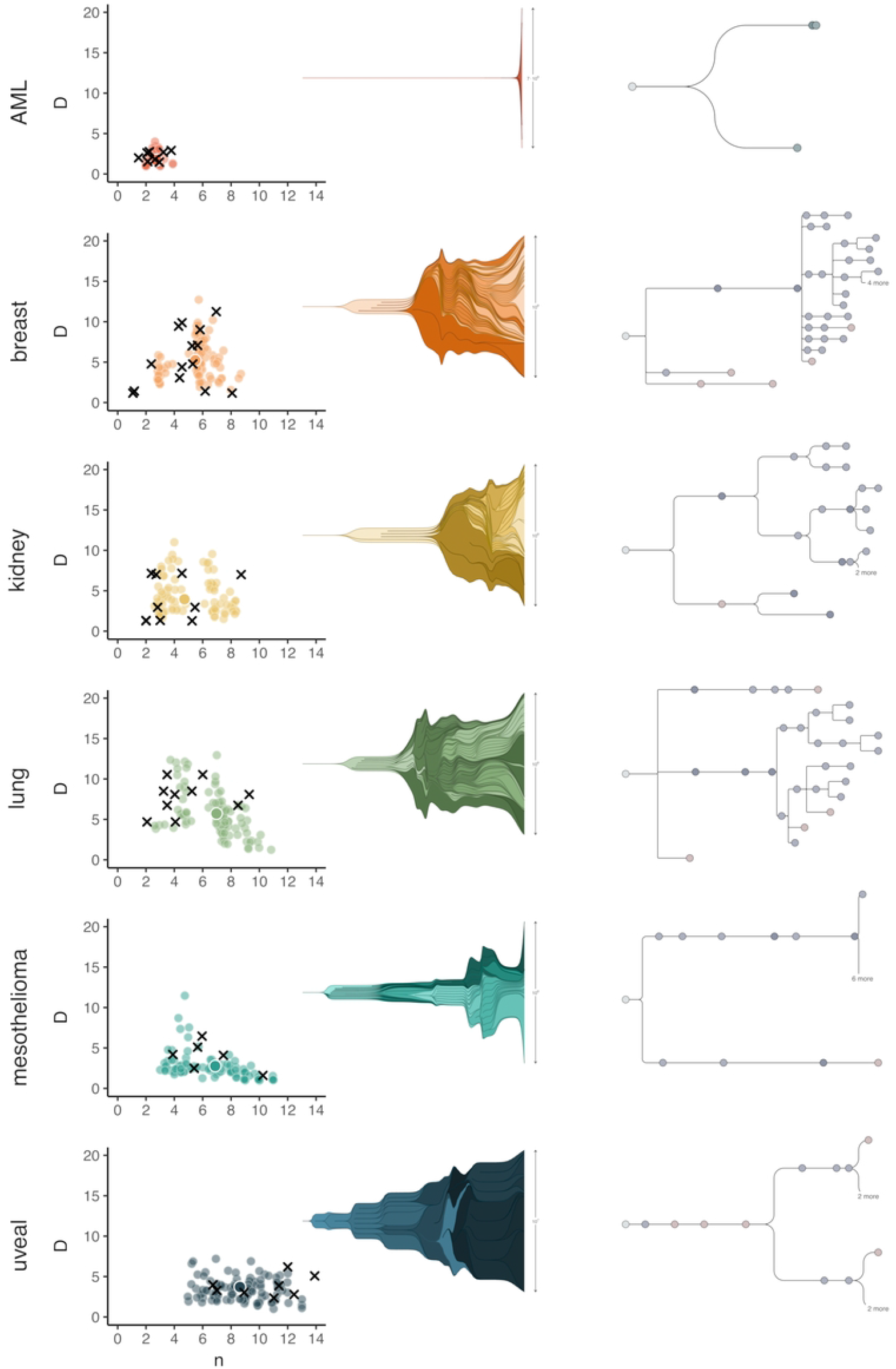

**Figure suppl S3_15.**
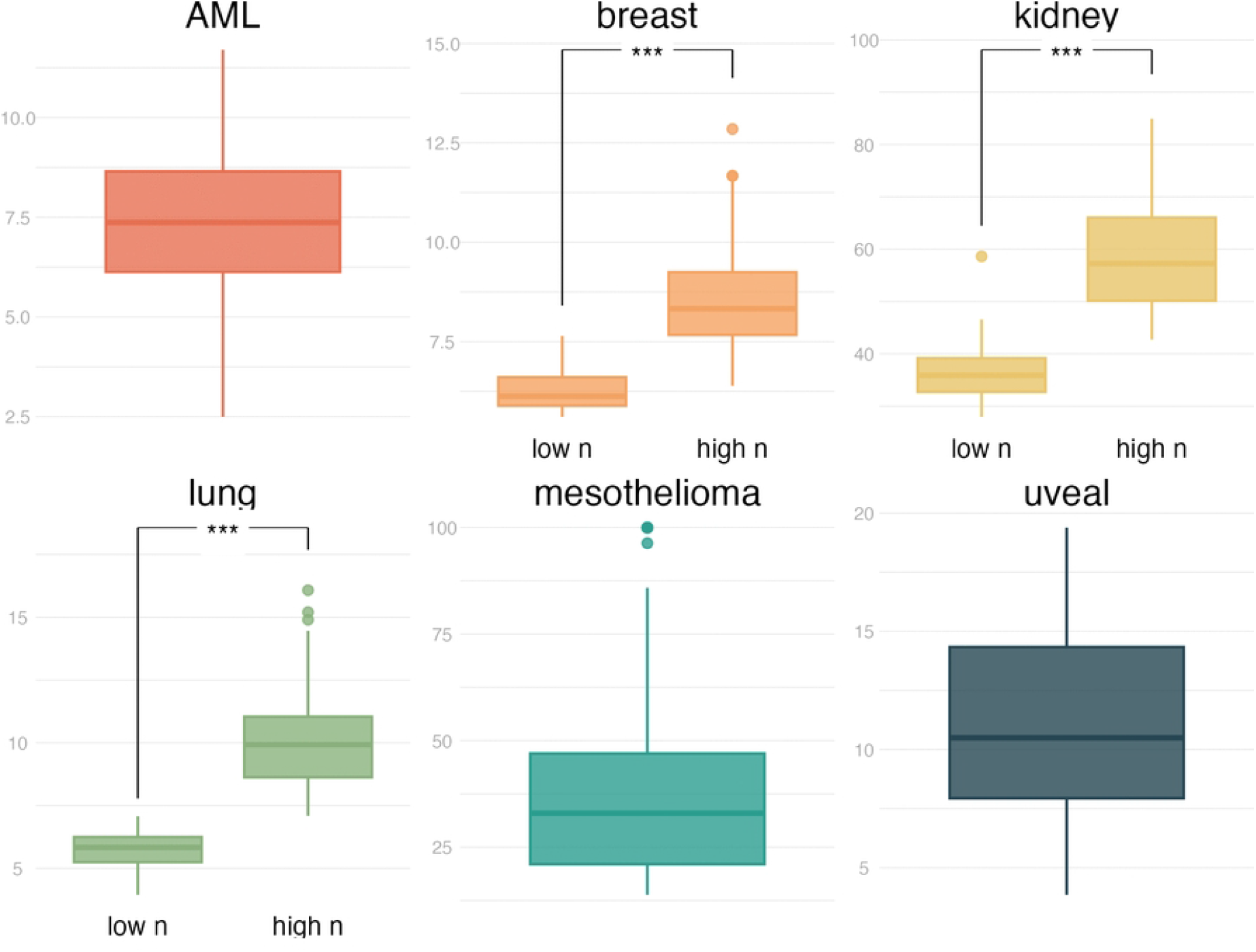

**Figure suppl S3_2.**
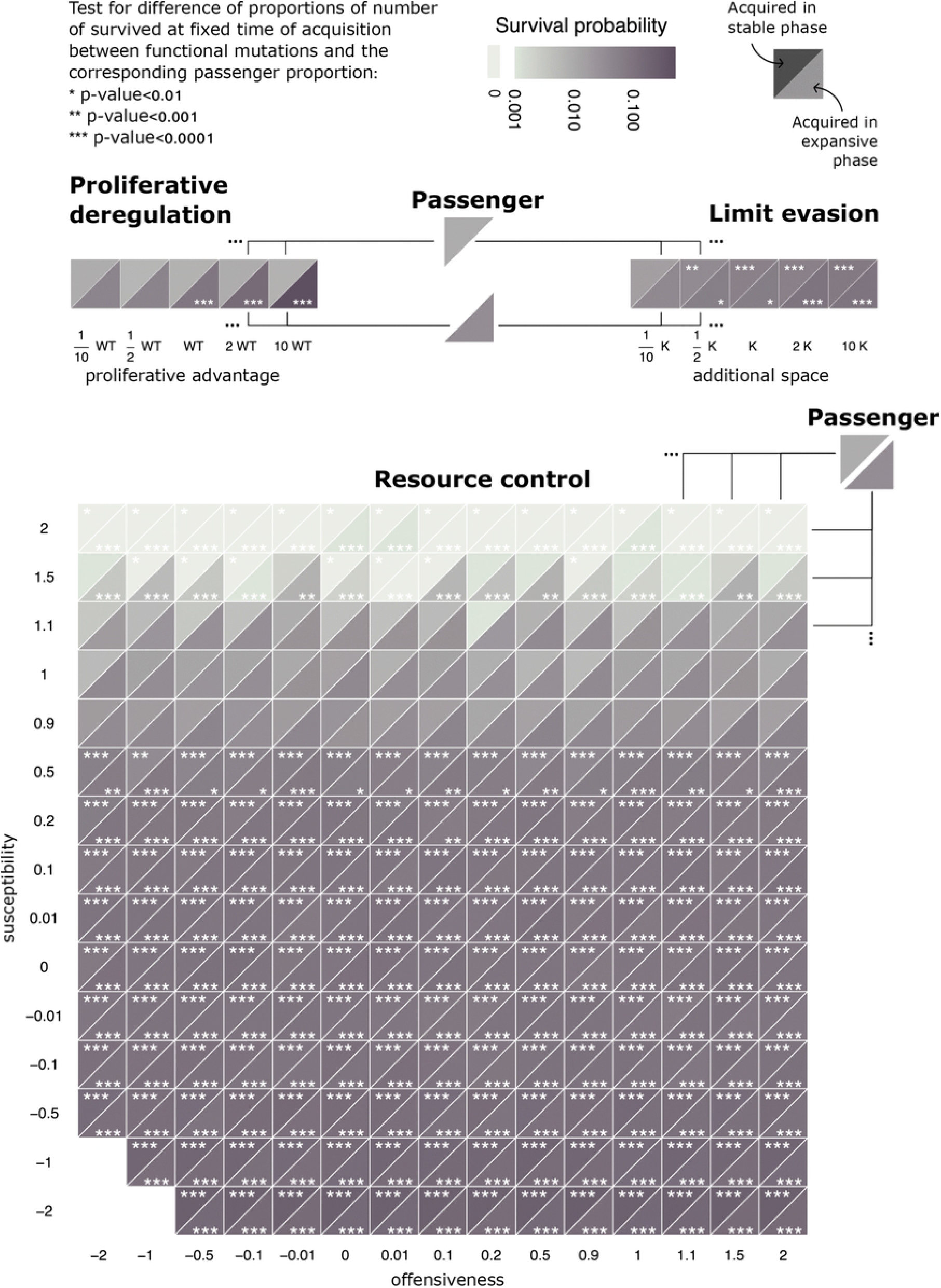

**Figure suppl S3_3.**
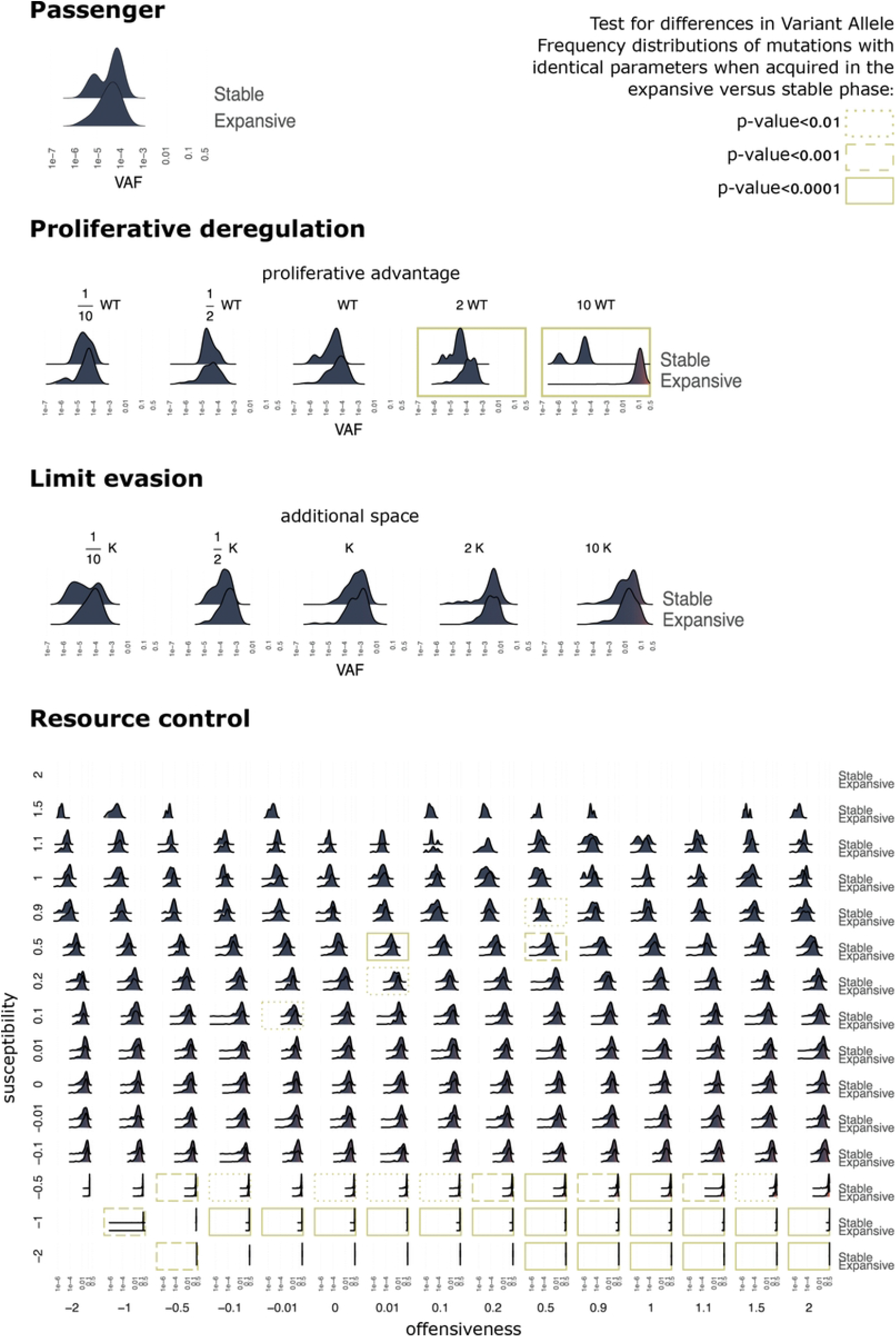

**Figure suppl S3_6.**
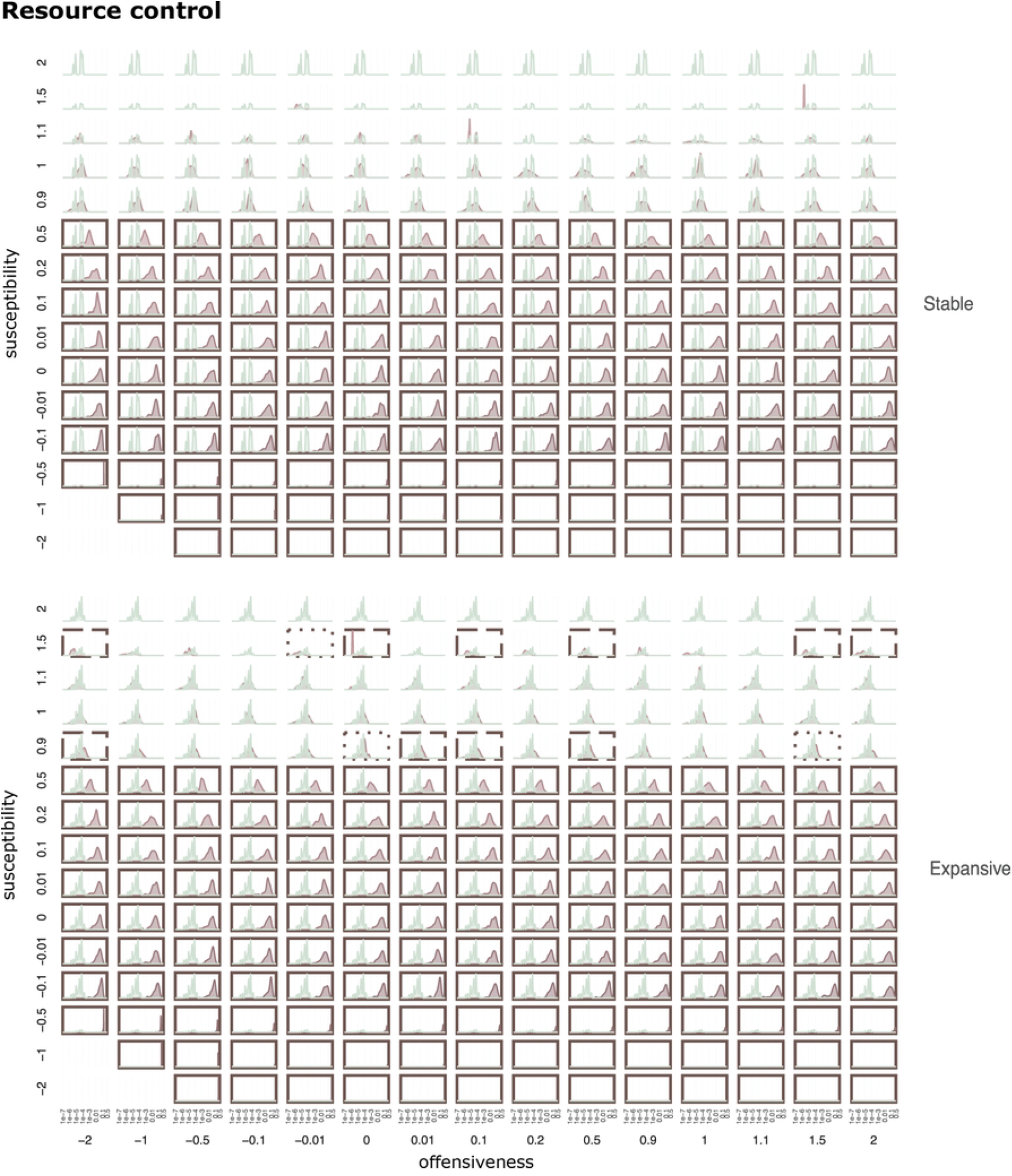

**Figure suppl S3_7.**
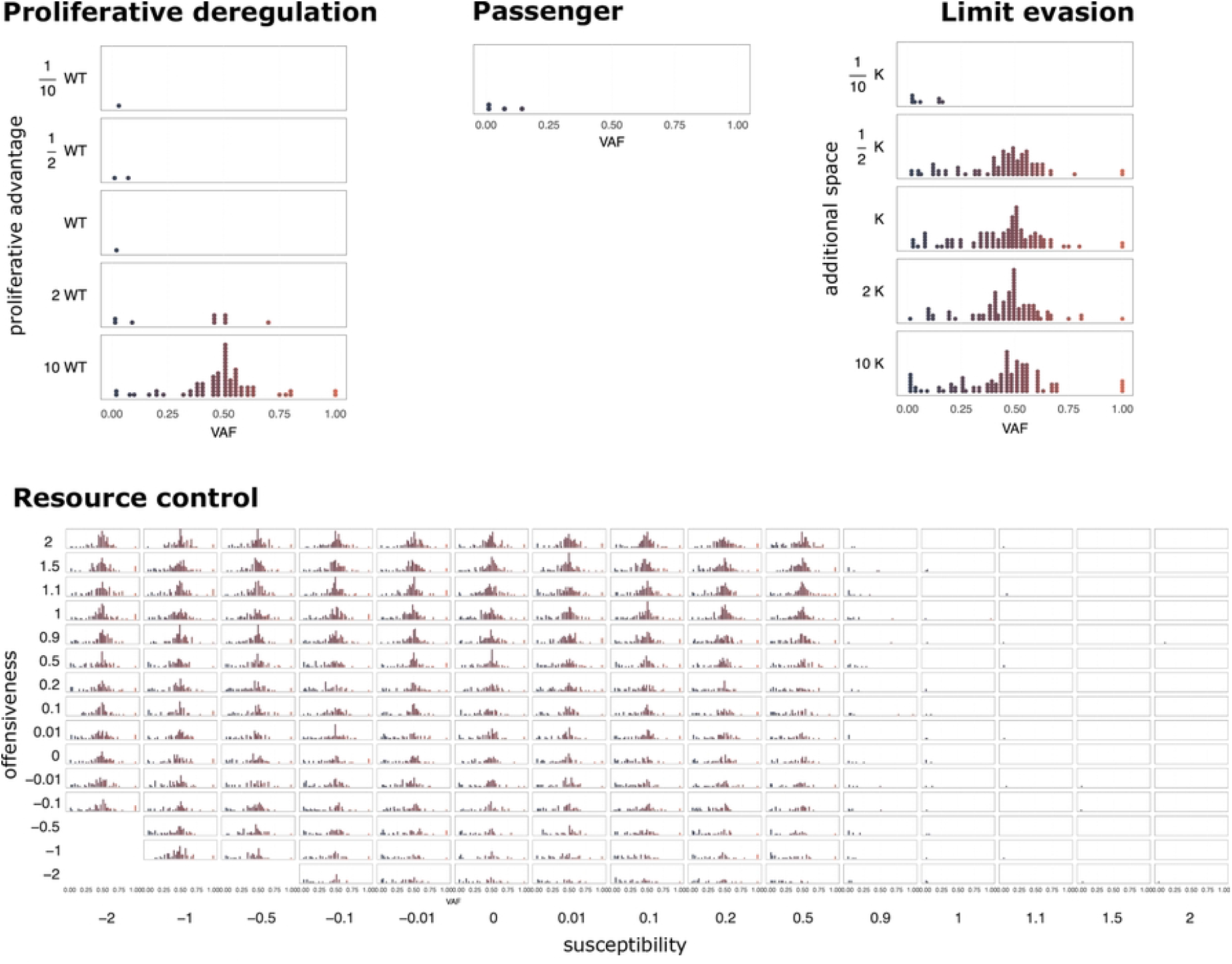

**Figure suppl S3_8.**
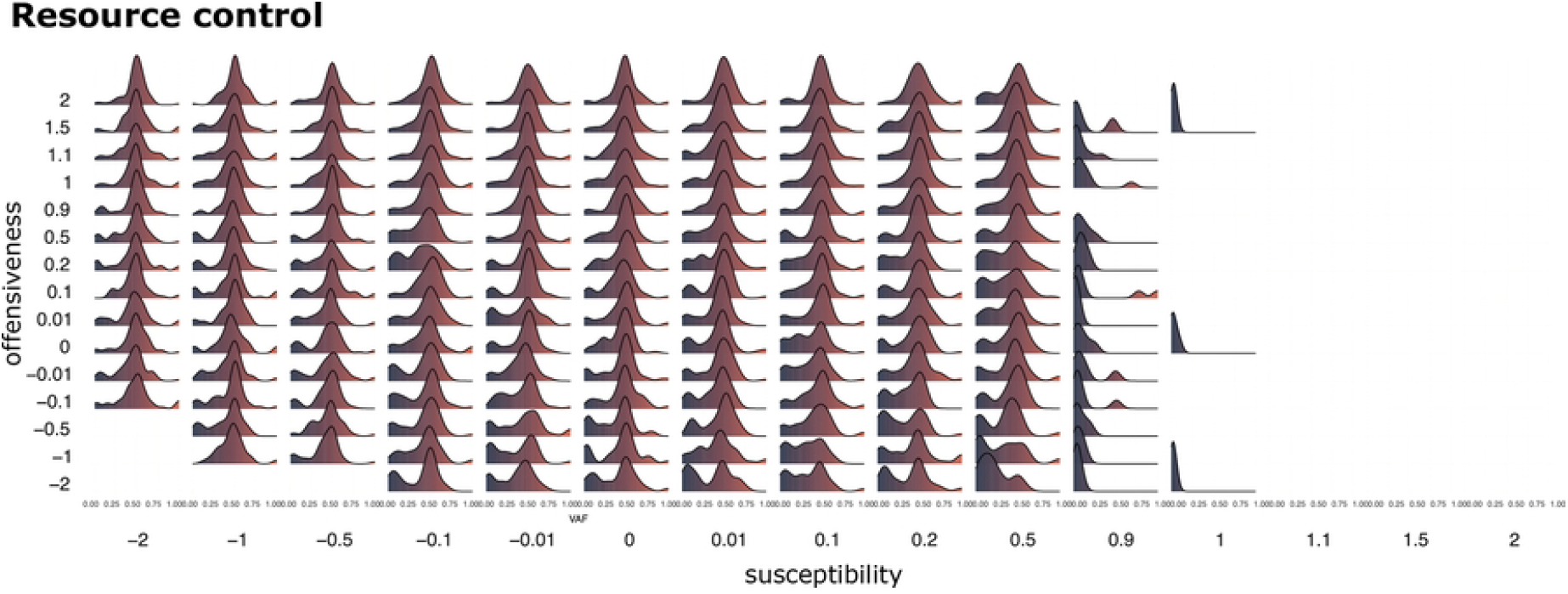

**Figure suppl S3_9.**
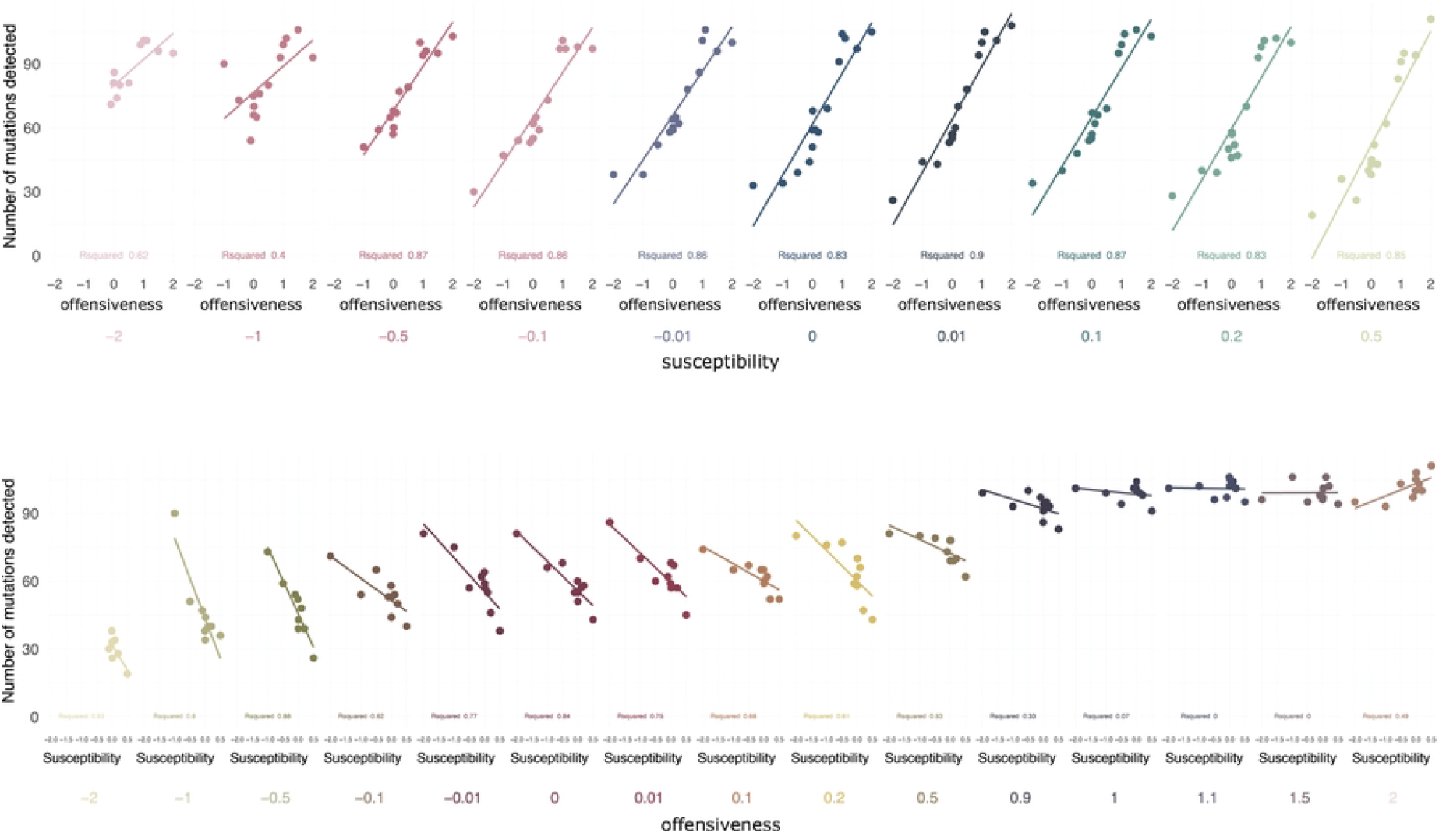

**Figure suppl S3_4.**
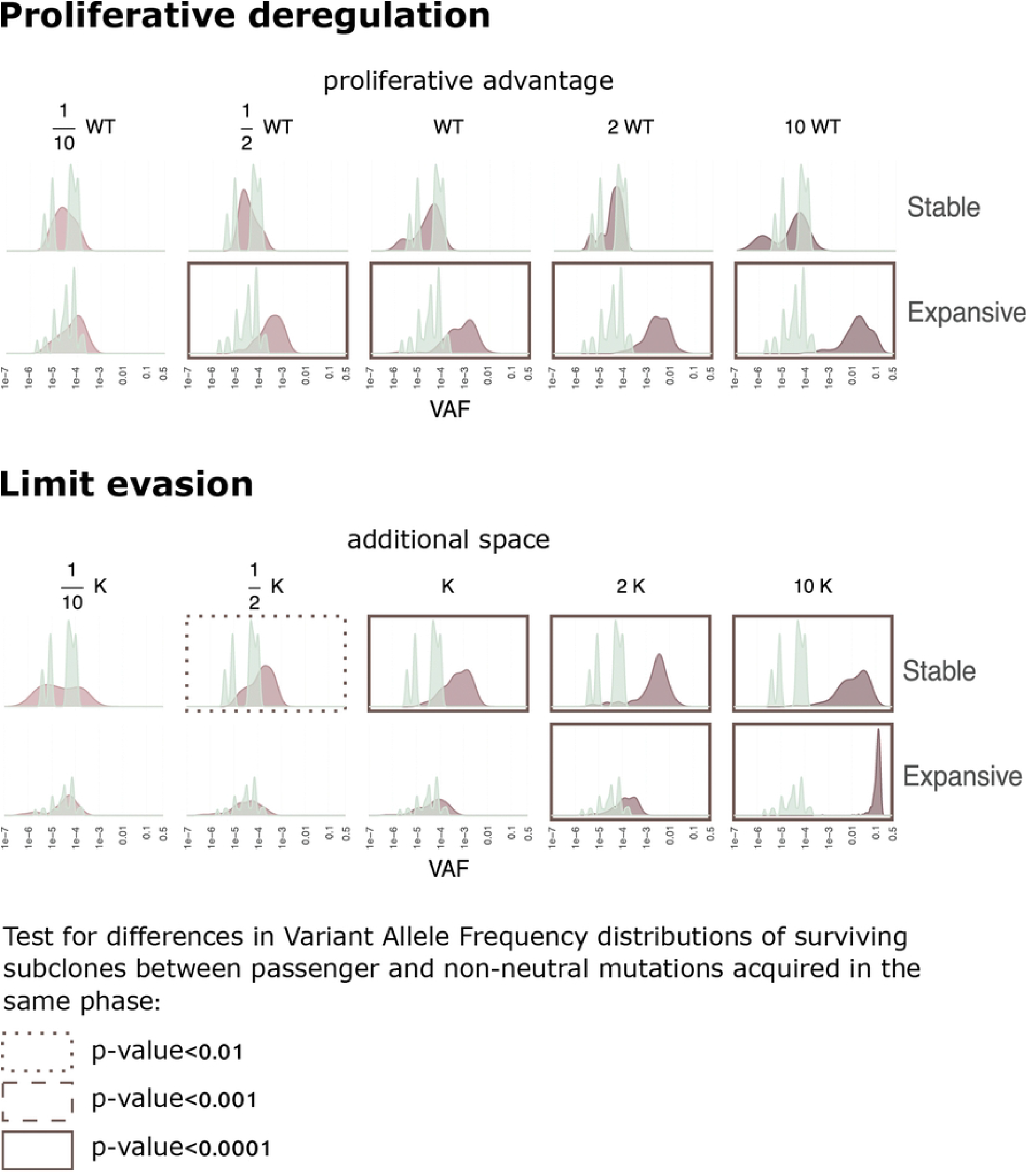

**Figure suppl S3_5.**
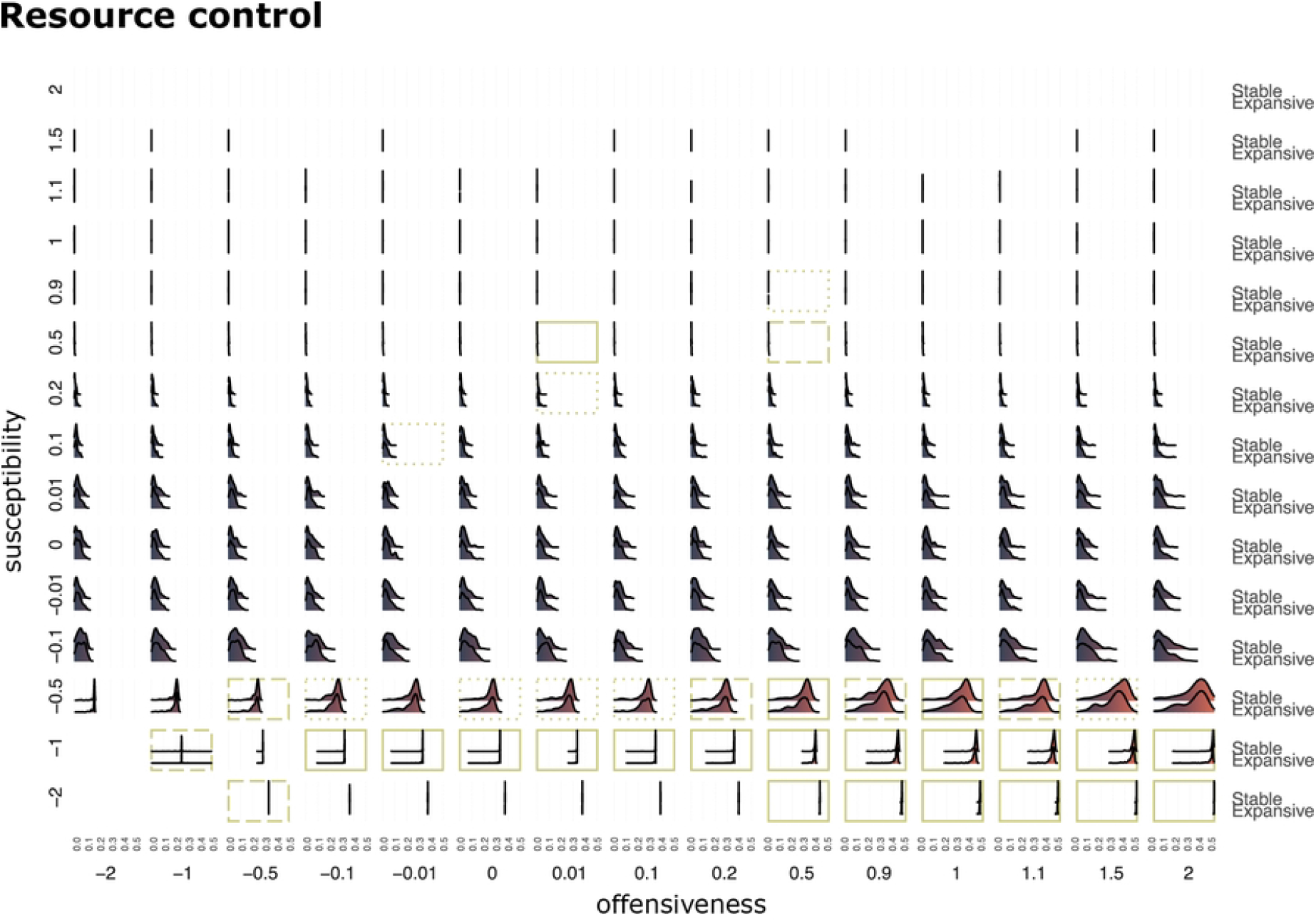

## References

1. Nowell PC. The Clonal Evolution of Tumor Cell Populations: Acquired genetic lability permits stepwise selection of variant sublines and underlies tumor progression. Science. 1976;194(4260):23–28. doi:10.1126/science.959840.

2. Marusyk A, Polyak K. Tumor heterogeneity: causes and consequences. Biochim Biophys Acta. 2010;1805(1):105–117.

3. Alizadeh AA, Aranda V, Bardelli A, Blanpain C, Bock C, Borowski C, et al. Toward understanding and exploiting tumor heterogeneity. Nature Medicine. 2015;21(8):846–853. doi:10.1038/nm.3915.

4. Michor F, Polyak K. The Origins and Implications of Intratumor Heterogeneity. Cancer Prevention Research. 2010;3(11):1361–1364. doi:10.1158/1940-6207.capr-10-0234.

5. Greaves M, Maley CC. Clonal evolution in cancer. Nature. 2012;481(7381):306–313. doi:10.1038/nature10762.

6. Burrell RA, McGranahan N, Bartek J, Swanton C. The causes and consequences of genetic heterogeneity in cancer evolution. Nature. 2013;501(7467):338–345. doi:10.1038/nature12625.

7. McGranahan N, Swanton C. Clonal Heterogeneity and Tumor Evolution: Past, Present, and the Future. Cell. 2017;168(4):613–628. doi:10.1016/j.cell.2017.01.018.

8. Burrell RA, Swanton C. Re-Evaluating Clonal Dominance in Cancer Evolution. Trends in Cancer. 2016;2(5):263–276. doi:10.1016/j.trecan.2016.04.002.

9. Mayers JR, Torrence ME, Danai LV, Papagiannakopoulos T, Davidson SM, Bauer MR, et al. Tissue of origin dictates branched-chain amino acid metabolism in mutantKras-driven cancers. Science. 2016;353(6304):1161–1165. doi:10.1126/science.aaf5171.

10. Davis A, Gao R, Navin N. Tumor evolution: Linear, branching, neutral or punctuated? Biochim Biophys Acta Rev Cancer. 2017;1867(2):151–161.

11. Merlo LMF, Pepper JW, Reid BJ, Maley CC. Cancer as an evolutionary and ecological process. Nature Reviews Cancer. 2006;6(12):924–935. doi:10.1038/nrc2013.

12. Altrock PM, Liu LL, Michor F. The mathematics of cancer: integrating quantitative models. Nature Reviews Cancer. 2015;15(12):730–745. doi:10.1038/nrc4029.

13. Beerenwinkel N, Schwarz RF, Gerstung M, Markowetz F. Cancer Evolution: Mathematical Models and Computational Inference. Systematic Biology. 2014;64(1):e1–e25. doi:10.1093/sysbio/syu081.

14. Fearon ER, Vogelstein B. A genetic model for colorectal tumorigenesis. Cell. 1990;61(5):759–767. doi:10.1016/0092-8674(90)90186-i.

15. Bozic I, Antal T, Ohtsuki H, Carter H, Kim D, Chen S, et al. Accumulation of driver and passenger mutations during tumor progression. Proceedings of the National Academy of Sciences. 2010;107(43):18545–18550. doi:10.1073/pnas.1010978107.

16. Gerlinger M, Rowan AJ, Horswell S, Larkin J, Endesfelder D, Gronroos E, et al. Intratumor Heterogeneity and Branched Evolution Revealed by Multiregion Sequencing. New England Journal of Medicine. 2012;366(10):883–892. doi:10.1056/nejmoa1113205.

17. Navin N, Kendall J, Troge J, Andrews P, Rodgers L, McIndoo J, et al. Tumour evolution inferred by single-cell sequencing. Nature. 2011;472(7341):90–94. doi:10.1038/nature09807.

18. Wang Y, Waters J, Leung ML, Unruh A, Roh W, Shi X, et al. Clonal evolution in breast cancer revealed by single nucleus genome sequencing. Nature. 2014;512(7513):155–160. doi:10.1038/nature13600.

19. Gerlinger M, Horswell S, Larkin J, Rowan AJ, Salm MP, Varela I, et al. Genomic architecture and evolution of clear cell renal cell carcinomas defined by multiregion sequencing. Nature Genetics. 2014;46(3):225–233. doi:10.1038/ng.2891.

20. Nik-Zainal S, Van Loo P, Wedge DC, Alexandrov LB, Greenman CD, Lau KW, et al. The Life History of 21 Breast Cancers. Cell. 2012;149(5):994–1007. doi:10.1016/j.cell.2012.04.023.

21. Shah SP, Roth A, Goya R, Oloumi A, Ha G, Zhao Y, et al. The clonal and mutational evolution spectrum of primary triple-negative breast cancers. Nature. 2012;486(7403):395–399. doi:10.1038/nature10933.

22. Ling S, Hu Z, Yang Z, Yang F, Li Y, Lin P, et al. Extremely high genetic diversity in a single tumor points to prevalence of non-Darwinian cell evolution. Proceedings of the National Academy of Sciences. 2015;112(47). doi:10.1073/pnas.1519556112.

23. Williams MJ, Werner B, Barnes CP, Graham TA, Sottoriva A. Identification of neutral tumor evolution across cancer types. Nature Genetics. 2016;48(3):238–244. doi:10.1038/ng.3489.

24. Sottoriva A, Kang H, Ma Z, Graham TA, Salomon MP, Zhao J, et al. A Big Bang model of human colorectal tumor growth. Nature Genetics. 2015;47(3):209–216. doi:10.1038/ng.3214.

25. Anderson ARA, Quaranta V. Integrative mathematical oncology. Nature Reviews Cancer. 2008;8(3):227–234. doi:10.1038/nrc2329.

26. Byrne HM. Dissecting cancer through mathematics: from the cell to the animal model. Nature Reviews Cancer. 2010;10(3):221–230. doi:10.1038/nrc2808.

27. Colson C, Whiting FJ, Baker AM, Graham TA. Mathematical modelling of cancer cell evolution and plasticity. Current Opinion in Cell Biology. 2025;95:102558. doi:10.1016/j.ceb.2025.102558.

28. Yin A, Moes DJAR, van Hasselt JGC, Swen JJ, Guchelaar H. A Review of Mathematical Models for Tumor Dynamics and Treatment Resistance Evolution of Solid Tumors. CPT: Pharmacometrics & Systems Pharmacology. 2019;8(10):720–737. doi:10.1002/psp4.12450.

29. Lee ND, Kaveh K, Bozic I. Clonal interactions in cancer: Integrating quantitative models with experimental and clinical data. Seminars in Cancer Biology. 2023;92:61–73. doi:10.1016/j.semcancer.2023.04.002.

30. Aguadé-Gorgorió G, Anderson ARA, Solé R. Modeling tumors as complex ecosystems. iScience. 2024;27(9):110699. doi:10.1016/j.isci.2024.110699.

31. Axelrod R, Axelrod DE, Pienta KJ. Evolution of cooperation among tumor cells. Proceedings of the National Academy of Sciences. 2006;103(36):13474–13479. doi:10.1073/pnas.0606053103.

32. Anderson ARA. A hybrid mathematical model of solid tumour invasion: the importance of cell adhesion. Mathematical Medicine and Biology: A Journal of the IMA. 2005;22(2):163–186. doi:10.1093/imammb/dqi005.

33. Bozic I, Allen B, Nowak MA. Dynamics of targeted cancer therapy. Trends in Molecular Medicine. 2012;18(6):311–316. doi:10.1016/j.molmed.2012.04.006.

34. Durrett R. Branching process models of cancer. Cham: Springer; 2015.

35. Durrett R, Foo J, Leder K, Mayberry J, Michor F. Intratumor Heterogeneity in Evolutionary Models of Tumor Progression. Genetics. 2011;188(2):461–477. doi:10.1534/genetics.110.125724.

36. Cheek D, Antal T. Mutation frequencies in a birth–death branching process. The Annals of Applied Probability. 2018;28(6). doi:10.1214/18-aap1413.

37. Kolev M. Mathematical modelling of the competition between tumors and immune system considering the role of the antibodies. Mathematical and Computer Modelling. 2003;37(11):1143–1152. doi:10.1016/s0895-7177(03)80018-3.

38. Noble R, Burri D, Le Sueur C, Lemant J, Viossat Y, Kather JN, et al. Spatial structure governs the mode of tumour evolution. Nature Ecology & Evolution. 2021;6(2):207–217. doi:10.1038/s41559-021-01615-9.

39. Streck A, Kaufmann TL, Schwarz RF. SMITH: spatially constrained stochastic model for simulation of intra-tumour heterogeneity. Bioinformatics. 2023;39(3). doi:10.1093/bioinformatics/btad102.

40. Hanahan D, Weinberg RA. The Hallmarks of Cancer. Cell. 2000;100(1):57–70. doi:10.1016/S0092-8674(00)81683-9.

41. Hanahan D, Weinberg RA. Hallmarks of cancer: the next generation. cell. 2011;144(5):646–674.

42. Hanahan D. Hallmarks of cancer: new dimensions. Cancer discovery. 2022;12(1):31–46.

43. Klebaner FC. On population-size-dependent branching processes. Advances in Applied Probability. 1984;16(1):30–55. doi:10.2307/1427223.

44. Jagers P, Klebaner FC. Population-size-dependent and age-dependent branching processes. Stochastic Processes and their Applications. 2000;87(2):235–254. doi:10.1016/s0304-4149(99)00111-8.

45. Lambert A. The Branching Process with Logistic Growth. The Annals of Applied Probability. 2005;15(2):1506–1535.

46. Daley DJ, Vere-Jones D. An introduction to the theory of point processes. 2nd ed. Probability and Its Applications. New York, NY: Springer; 2003.

47. Bailey NT. The elements of stochastic processes with applications to the natural sciences. John Wiley & Sons; 1991.

48. Athreya P Krishna; Ney. Branching Processes. Berlin, Heidelberg: Springer Berlin Heidelberg; 1972.

49. Gillespie DT. Approximate accelerated stochastic simulation of chemically reacting systems. J Chem Phys. 2001;115(4):1716–1733.

50. Balaparya A, De S. Revisiting signatures of neutral tumor evolution in the light of complexity of cancer genomic data. Nature Genetics. 2018;50(12):1626–1628. doi:10.1038/s41588-018-0219-4.

51. McDonald TO, Chakrabarti S, Michor F. Currently available bulk sequencing data do not necessarily support a model of neutral tumor evolution. Nature Genetics. 2018;50(12):1620–1623. doi:10.1038/s41588-018-0217-6.

52. Tarabichi M, Martincorena I, Gerstung M, Leroi AM, Markowetz F, Spellman PT, et al. Neutral tumor evolution? Nature Genetics. 2018;50(12):1630–1633. doi:10.1038/s41588-018-0258-x.

53. Morita K, Wang F, Jahn K, Hu T, Tanaka T, Sasaki Y, et al. Clonal evolution of acute myeloid leukemia revealed by high-throughput single-cell genomics. Nature Communications. 2020;11(1). doi:10.1038/s41467-020-19119-8.

54. Turajlic S, Xu H, Litchfield K, Rowan A, Chambers T, Lopez JI, et al. Tracking Cancer Evolution Reveals Constrained Routes to Metastases: TRACERx Renal. Cell. 2018;173(3):581–594.e12. doi:10.1016/j.cell.2018.03.057.

55. Zhang M, Luo JL, Sun Q, Harber J, Dawson AG, Nakas A, et al. Clonal architecture in mesothelioma is prognostic and shapes the tumour microenvironment. Nature Communications. 2021;12(1). doi:10.1038/s41467-021-21798-w.

56. Minussi DC, Nicholson MD, Ye H, Davis A, Wang K, Baker T, et al. Breast tumours maintain a reservoir of subclonal diversity during expansion. Nature. 2021;592(7853):302–308. doi:10.1038/s41586-021-03357-x.

57. Yates LR, Gerstung M, Knappskog S, Desmedt C, Gundem G, Van Loo P, et al. Subclonal diversification of primary breast cancer revealed by multiregion sequencing. Nature Medicine. 2015;21(7):751–759. doi:10.1038/nm.3886.

58. Jamal-Hanjani M, Wilson GA, McGranahan N, Birkbak NJ, Watkins TBK, Veeriah S, et al. Tracking the Evolution of Non–Small-Cell Lung Cancer. New England Journal of Medicine. 2017;376(22):2109–2121. doi:10.1056/nejmoa1616288.

59. Durante MA, Rodriguez DA, Kurtenbach S, Kuznetsov JN, Sanchez MI, Decatur CL, et al. Single-cell analysis reveals new evolutionary complexity in uveal melanoma. Nature Communications. 2020;11(1). doi:10.1038/s41467-019-14256-1.

60. Milholland B, Dong X, Zhang L, Hao X, Suh Y, Vijg J. Differences between germline and somatic mutation rates in humans and mice. Nature Communications. 2017;8(1). doi:10.1038/ncomms15183.

61. Del Monte U. Does the cell number 109still really fit one gram of tumor tissue? Cell Cycle. 2009;8(3):505–506. doi:10.4161/cc.8.3.7608.

62. Janssen WJ, Bratton DL, Jakubzick CV, Henson PM. Myeloid Cell Turnover and Clearance. Microbiology Spectrum. 2016;4(6). doi:10.1128/microbiolspec.mchd-0005-2015.

63. Bowden DH. Cell turnover in the lung. The American Review of Respiratory Disease. 1983;128(2 Pt 2):S46–S48. doi:10.1164/arrd.1983.128.2P2.S46.

64. Ksiek K. Mesothelial cell: A multifaceted model of aging. Ageing Research Reviews. 2013;12(2):595–604. doi:10.1016/j.arr.2013.01.008.

65. Chen J, Zhang H, Yi X, Dou Q, Yang X, He Y, et al. Cellular senescence of renal tubular epithelial cells in acute kidney injury. Cell Death Discovery. 2024;10(1). doi:10.1038/s41420-024-01831-9.

66. Hu D. Regulation of Growth and Melanogenesis of Uveal Melanocytes. Pigment Cell Research. 2000;13(8):81–86. doi:10.1034/j.1600-0749.13.s8.15.x.

67. for Disease Control C, Program PWTCH. Minimum Latency & Types or Categories of Cancer. U.S. Department of Health and Human Services; 2015. Available from: https://www.cdc.gov/wtc/pdfs/policies/WTCHP-Minimum-Cancer-Latency-PP-01062015-508.pdf.

68. Abecasis M, Cross NCP, Brito M, Ferreira I, Sakamoto KM, Hijiya N, et al. Is cancer latency an outdated concept? Lessons from chronic myeloid leukemia. Leukemia. 2020;34(9):2279–2284. doi:10.1038/s41375-020-0957-z.

69. Ornstein MC, Mukherjee S, Mohan S, Elson P, Tiu RV, Saunthararajah Y, et al. Predictive factors for latency period and a prognostic model for survival in patients with therapy-related acute myeloid leukemia. American Journal of Hematology. 2013;89(2):168–173. doi:10.1002/ajh.23605.

70. Olsson H, Baldetorp B, Fernö M, Perfekt R. Relation between the rate of tumour cell proliferation and latency time in radiation associated breast cancer. BMC Cancer. 2003;3(1). doi:10.1186/1471-2407-3-11.

71. Molinari L. Mesothelioma Latency Period; n.d. Available from: https://www.mesothelioma.com/mesothelioma/latency-period/.

